# Deep sequence-function mapping reveals computationally unpredictable epistatic trajectories that drive laboratory evolution

**DOI:** 10.64898/2026.11.11.766033

**Authors:** F. E. H. Nintzel, M. Lihan, M. Penner, M. Sieders, M. Gantz, H. Bormke, C. Sifri, E. J. Medcalf, S. V. Mathis, S. Cao, F. Hollfelder

## Abstract

The black-box nature of directed evolution has been constraining advances in enzyme engineering and the wider application of engineered biocatalysts. While conventional deep sequence-function mapping has delivered insights into sequence space architecture, it lacks the ability to analyse adaptive landscapes and fully grasp the long-term impact of epistasis on the accessibility of evolutionary trajectories. Here, we introduce Deep Evolutionary Trajectory Mapping (dETM) which tracks the winners of a multi-round evolutionary campaign to map catalytic rates across the adaptive landscape of an imine reductase. We identified founder variants as stepping stones towards fitness peaks and recognise positive epistasis as their competitive advantage over alternative lineages. State-of-the-art machine learning models could not predict the resulting winning trajectories, which highlights the importance of decoding design principles for successful evolution experimentally. Permissive selection and ultrahigh-throughput screening emerge as foundations of derisking landscape traversal by keeping multiple paths accessible and avoiding diminishing returns.

## Introduction

Directed evolution is widely used to improve proteins for new functions of interest (*1*), yet its outcomes remain difficult to predict. Ultrahigh throughput screening technologies make a productive outcome more likely, but the sheer number of possible amino acid combinations means that success is not guaranteed. As long as the fundamental mechanisms that drive evolutionary outcomes remain elusive, directed evolution will remain cumbersome, with labour-intensive hit-and-miss campaigns at the mercy of chance. To avoid wasting substantial experimental capacity on marginal advances in multi-round campaigns, a framework for deriving the rules of sequence space navigation is necessary.

The difficulty of traversing an enzyme’s fitness landscape with directed evolution lies in the complex architecture of sequence space. Beneficial mutations are rare and paths to improving enzyme fitness can be disrupted by non-linear interactions between mutations (*2–4*). Such non-linearity arises from intra-gene epistasis, whereby the fitness effect of a mutation depends on the genetic background in which it occurs (*5*). Negative epistasis can render combinations of individually beneficial mutations less advantageous than expected. A preponderance of negatively epistatic interactions can constrain evolution by blocking multi-step mutational pathways towards optimal function (*6–8*), causing diminishing returns (*9*) or creating evolutionary dead-ends (*10*).

With the advent of high throughput screening and deep sequencing, molecular evolution and enzymology have begun to transform into -omics disciplines (*11*). Progress towards large datasets that can be quantitatively evaluated has benefited from deep mutational scanning (DMS), which combines large-scale mutagenesis, high-throughput screening and deep sequencing to evaluate mutational effects on protein properties (*12*). DMS datasets have been used to answer fundamental questions about the architecture of sequence space (*13*), study the evolvability of individual enzymes (*14*), and inform engineering campaigns through single rounds of AI-extrapolation (*15*) or multiple active-learning cycles (*16*).

Deep mutational scanning typically maps mutational effects around a defined genetic background and therefore provides a largely static and local view of a fitness landscape. Consequently, it misses how the landscape is reshaped as evolution proceeds and does not reveal the ‘forward network’ of possible mutations that determines which trajectories remain accessible across changing genetic backgrounds (*17*). Multi-round evolution is therefore required to explore how epistatic networks change during adaptive evolution and shape the accessibility of evolutionary trajectories (*18*). Multi-round campaigns also permit dissection of how mutational effects depend on genetic starting points (*5, 17, 19*), the timing of population-level events (*8, 20*) and evolutionary paths previously taken (*21, 22*).

A vast record of multi-round evolution data is encoded in the sequence diversity of natural proteins. Multiple sequence alignments have long been used to leverage this evolutionary information to infer constraints on protein sequence-function relationships (*23, 24*). More recently, unsupervised protein language models have used evolutionary sequence information to predict beneficial mutations and guide trajectories towards fitness peaks (*25, 26*). However, it remains unclear whether evolutionary history can unambiguously predict the epistatic trajectories that become accessible across changing genetic backgrounds during laboratory evolution, so collection of dynamic datasets in response to a well defined fitness challenge become attractive.

Here, we introduce Deep Evolutionary Trajectory Mapping (dETM), a framework based on collecting a variant library from a multi-round directed evolution campaign and labelling each collected variant with quantitative fitness scores reporting on enzymatic rates. The resulting dataset enables us to investigate how adaptive trajectories emerge across changing genetic backgrounds. We applied dETM to five rounds of directed evolution of an imine reductase, using error-prone PCR to generate predominantly single-step mutational diversity and droplet microfluidics for ultrahigh-throughput screening (*27*). We further compared the observed adaptive trajectories with natural evolutionary sequence information and predictions from unsupervised protein-language models to determine whether the paths taken during laboratory evolution could have been predicted computationally. We found that the population became organised around a small number of founder variants which benefited from positive epistasis and reshaped mutational effects across subsequent rounds: constructive trajectories were directed by founder-compatible mutations along towards the campaign’s fitness peak, while previously beneficial but incompatible mutations became evolutionary dead ends. The resulting winning trajectory deviated from the natural sequence record and none of the tested computational models predicted productive steps across the campaign better than a random baseline. These limits to current computational prediction reinforce the value of experimental evolution and underline the need for design principles that guide the process of evolution towards a successful, unprecedented outcome. Here, combining permissive selection with ultrahigh-throughput screening allowed the luxury to invest exploration effort in variants with modest immediate fitness returns but the potential for much larger future payoffs through positive epistasis - a patient walk through sequence space.

### Dissecting adaptive evolution with Deep Evolutionary Trajectory Mapping

In this study, we have examined the adaptive laboratory evolution of the bacterial imine reductase *Sr*IRED (*28, 29*) which catalyses the formation of chiral amine bonds from carbonyl and amine substrates. We used this system to model a common adaptive evolutionary format over five rounds in which variant libraries were generated using error-prone PCR and screened for improved catalytic activity towards the model substrates cyclohexanone **1** and cyclopropylamine **a** (**Figure 1A and Figure S1)**. We used absorbance-activated droplet sorting (AADS, **Figure S2)** (*30*) for microfluidics-based ultra-high throughput screening of libraries comprising ∼200,000 unique variants per round (**Figure 1C and Figure S1)**. Enzymatic turnover was quantified by colorimetric detection of the WST-1 formazan dye generated from the unconverted cofactor remaining after the IRED reaction (*15*). Each round, the most active 200 - 500 variants were selected and carried forward as template for the next round of diversification (**Figure 1A** and **Figure S1**). Screening success was validated by plate-based enzymatic assays of cell lysates from 90 randomly selected variants (**Figure S3**). Across five rounds of evolution, we screened approximately one million mutants and selected the 1,342 most active variants (**Figure 1A** and **Figure S1**).

**Figure 1:**
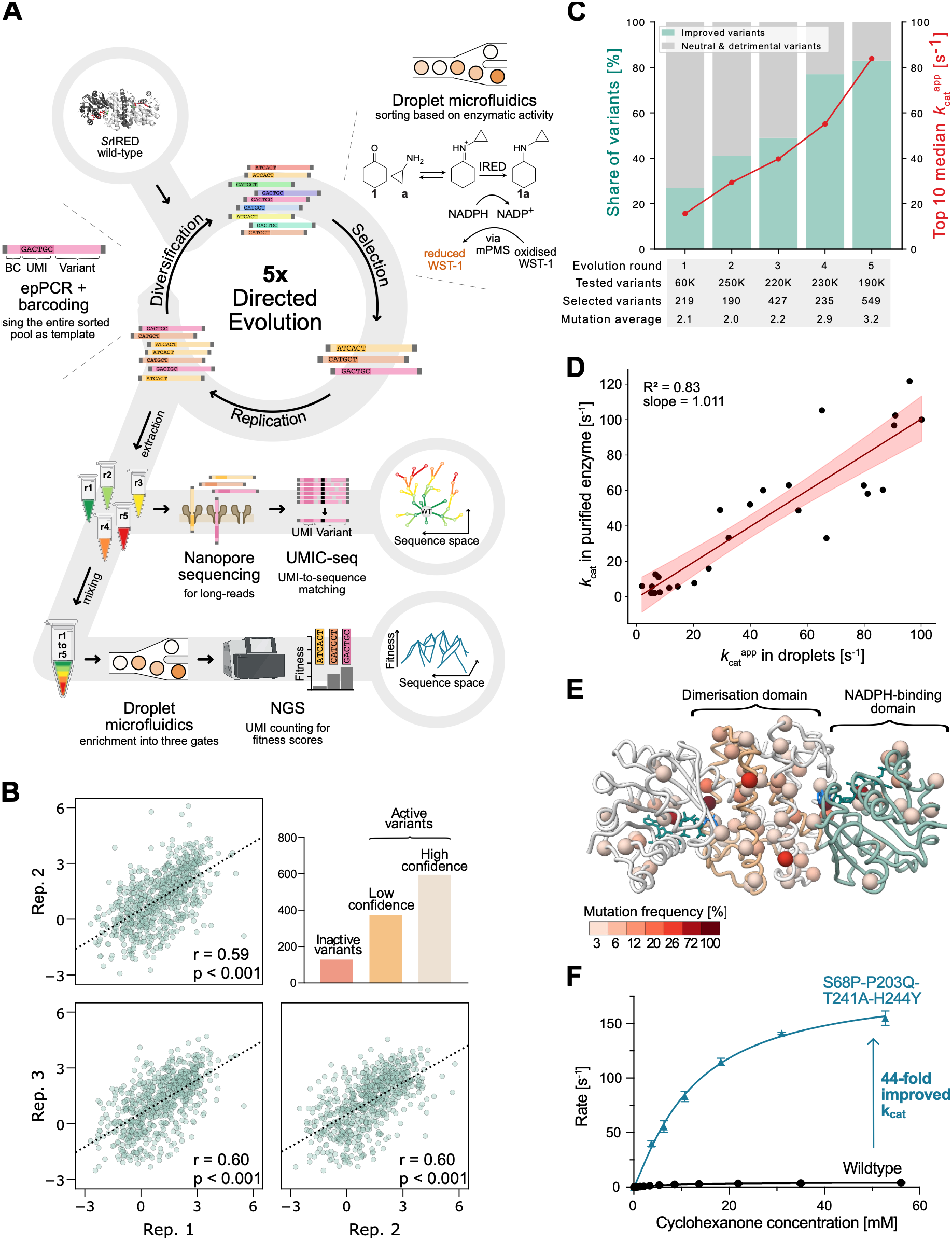
Recording the adaptive evolution of an IRED with microfluidics and deep sequencing. **(A)** Deep sequence-function mapping workflow consisting of (i) five rounds of library generation of unique molecular identifier (UMI)- and barcode (BC)-tagged variants, (ii) screening in microdroplets against substrates cyclohexanone **1** and cyclopropylamine **a** and (iii) using the entire pool of sorted variants to start the next round of library generation and selection with increased stringency. Long-read nanopore sequence was used to derive UMI-variant linkage for the sorted pool of each round. The fitness scores of the entire sorted pool from the campaign were recorded with a microdroplet-based enrichment experiment in triplicate using the short UMIs for NGS enrichment quantification. **(B)** Correlation between high-confidence enrichment score replicates. A total of 1071 variants with enrichment score were recovered, among them 594 variants that carried a high confidence label. **(C)** Overview of the directed evolution campaign using the high-confidence dataset. The fraction of improving variants in the selected pool (+1 σ over WT fitness) as well as the median *k* ^app^ of the top 10 variants per round increased constantly over the course of the directed evolution campaign. **(D)** Correlation between experimentally derived *k*_cat_ values of purified proteins and *k* ^app^ values based on the exponential fit of enrichment scores. **(E)** Structural mapping (PDB 5ocm) of mutations present in the 36 hits (variants with *k* ^app^ 4-fold over WT) after five rounds of directed evolution. Mutations accumulated during evolution are widely distributed over the structure with higher mutation frequency at the dimer interface (orange) and the substrate and cofactor binding domains (green). **(F)** Michaelis-Menten kinetics of *Sr*IRED wildtype and the campaign’s winner S68P-T241A-P203Q-H244Y reveal a 44-fold improvement in *k*_cat_ from 4.7 ± 0.1 s^-1^ for wildtype to 205.2 ± 5.5 s^-1^ for the quadruple variant.

To investigate how adaptive trajectories emerged during this evolution campaign, we established Deep Evolutionary Trajectory Mapping (dETM), an approach that combines variant identity and quantitative fitness measurements for the winning variants. We used long-read UMI-linked consensus sequencing (UMIC-seq) with the Oxford Nanopore technology (*31*) to capture the identity of the 1,342 winners and to resolve mutational co-occurrence of amino acids for epistasis analysis (**Figure 1A**). To quantify the fitness of the selected variants, we pooled all 1,342 winners into a composite library while preserving their round-specific barcodes. The pooled library was screened in triplicates using the above-described microfluidic workflow into high-, low- and no-activity bins. NGS was used to quantify the occurrence of individual UMIs in the input library as well as in the high- and low-activity output bins, with UMI counts mapped to variant identities using the UMIC-seq dictionary (**Figure 1A, Figure S4** and **Figure S5**). The final dETM dataset contained 1071 highly replicable fitness labels for the winners of five rounds of directed evolution (r = 0.6 and p < 0.001, **Figure 1B, Figure S1 and Figure S6**).

Correlating our droplet-based fitness scores with biophysical and kinetic parameters (T_M_, *k*_cat_, K_M_, *k*_cat_/K_M_) of 27 purified enzyme variants revealed the primary evolutionary driving force of this campaign to have been catalytic rate (*k*_cat_, **Figure S7**). Thus, we translated our fitness scores into *k* ^app^ labels for the 1071 observed variants in the dETM dataset (R^2^ = 0.83, **Figure 1D**). The strong correlation between fitness and catalytic rate enabled mechanistic interpretation of the evolutionary data which was previously not possible with conventional fitness scores representing a composite of multiple protein traits (i.e. stability, expression level, catalytic rate, and substrate affinity) (*32*).

The dETM dataset revealed a population-wide improvement in *k*_cat_^app^ over the course of the campaign (**Figure S3**). The share of improved variants (+1 SD over wildtype *k*_cat_^app^) increased consistently round-on-round from 27% in round 1 to 84% in round 5, indicating that the population evolved without fitness collapse (**Figure 1C**). In addition, the quality of hits generated during the campaign (quantified by the median *k*_cat_^app^ of the top 10 variants per round) increased consistently over the five rounds (**Figure 1C, Table S1**). Structural mapping of the positions which were mutated in the top hits highlighted that mutations were widely spread over the structure with many mutations occurring in the dimerization domain as well as the NADPH-binding domain (**Figure 1E, Figure S8**). We identified the quadruple mutant S68P-T241A-P203Q-H244Y as the fitness peak of the campaign with Michaelis-Menten kinetics of the purified enzyme confirming a 44-fold increase in *k*_cat_ (205 ± 5.5 s^-1^ *vs.* 4.7 ± 0.1 s^-1^ for wildtype, **Figure 1F**, **Table S2** and **Figure S9**).

### Founder variants drive the exploration of sequence space

More than simply providing a catalogue of hits from the evolution campaign, the dETM dataset harbours the sequence-function relationships required to reconstruct the adaptive trajectories that produced the hits. We therefore represented the campaign as a network graph of sequence-space exploration (*33–35*). Each variant was represented as a node with edges connecting variants differing by one amino acid mutation (**Figure 2A** and **Figure S10**). We identified 144 trajectories leading to the campaign’s 36 unique hits. While most hits were reached by an average of two trajectories, a cluster of ten hits was accessed by an average of eight trajectories per hit. This highly connected cluster included variant S68P-T241A-P203Q-H244Y which was identified as fitness peak of the campaign (**Figure 2A)**.

**Figure 2:**
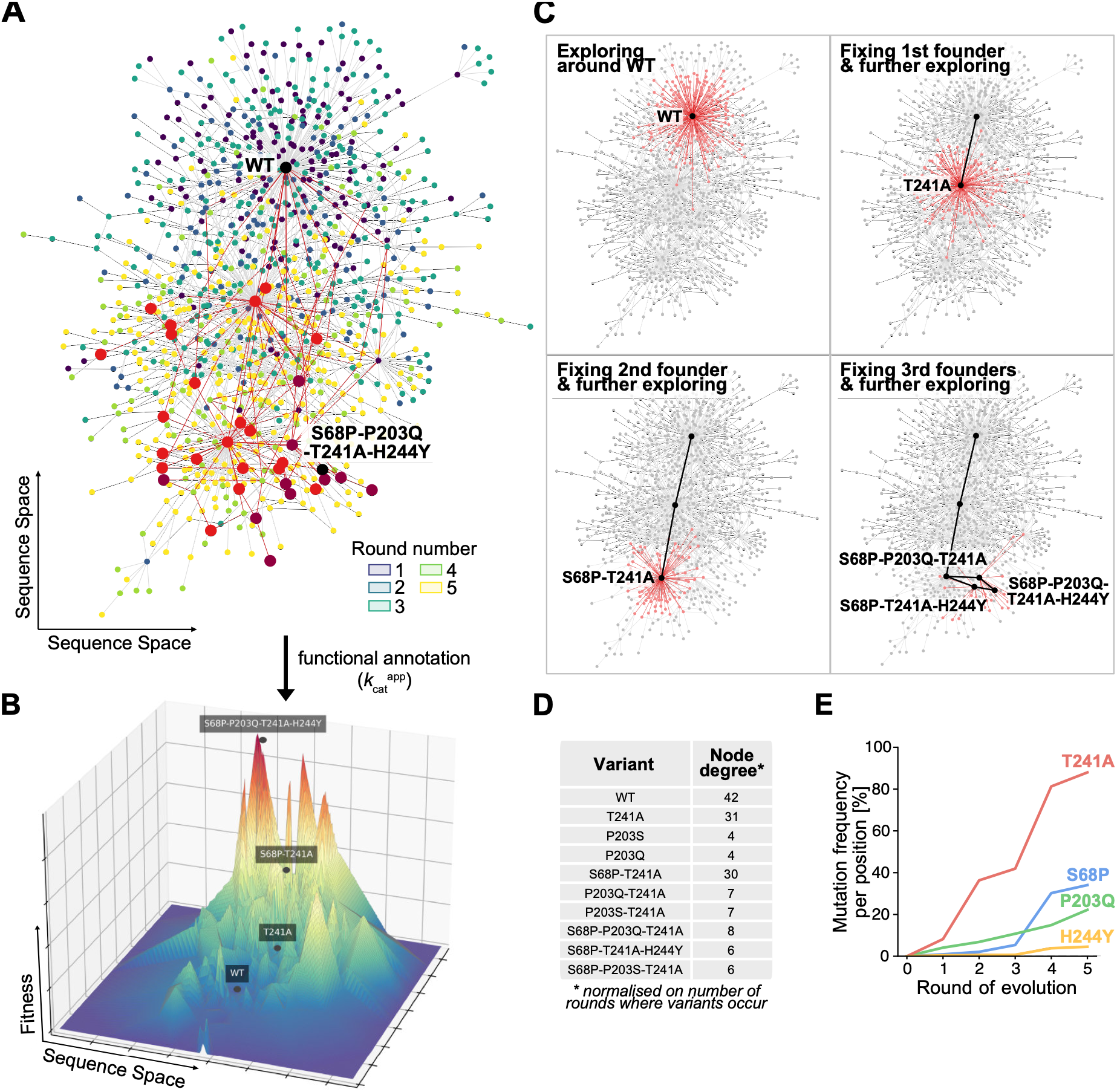
The exploration of sequence space and emergence of founder variants. **(A)** Network graph of the recorded evolution campaign where each node represents one variant (coloured by the first-occurrence round) and edges are connecting variants differing by one amino acid mutation. Wildtype (WT) and the campaign’s fitness peak (S68P-P203Q-T241A-H244Y) are marked in black. Hits of the directed evolution campaign (defined here as variants four-fold better than wild-type) and all possible evolutionary trajectories between wildtype and hits are marked in red. Those hits with the highest number of trajectories explored from wildtype (> 5 trajectories) are highlighted in dark red. **(B)** The traversed fitness landscape of the evolution campaign. A network graph of the variants with high-confidence fitness labels was used as x-y-plane and respective *k*_cat_^app^ scores were added as z-axis. Wildtype (WT), relevant intermediate mutations (T241A, S68P-T241A), and the campaign’s fitness peak (S68P-P203Q-T241A-H244Y) are highlighted. **(C)** Time-resolved representation of the recorded dETM data. Starting at wildtype, the campaign was replayed in single-mutation steps with variant progeny in red. Evolution followed an explore-and-fix pattern with stepwise exploration of sequence space followed by fixation of successful variants as founder variants. **(D)** Table of variants with highest node degrees, i.e. largest number of descendants per round. Node degrees were normalised by the number of rounds in which a variant occurred. Founder variants can be identified as variants with highest node degrees. **(E)** Time-resolved frequency at which a specific position in *Sr*IRED has undergone mutation across the entire assayed population. Founder mutations can be identified as predominant at some stage during the evolution campaign with frequencies >7%.

We interpret the number of pathways as an indicator of evolutionary accessibility, with each trajectory representing a solution for navigating epistatic constrains. The high number of trajectories per hit therefore suggests an evolvable starting point, where small mutational steps provided sufficient improving diversity to avoid local fitness maxima and evolutionary dead-ends (*6, 7, 36*). This interpretation was further corroborated when we enriched the network graph representation of the explored sequence space with our recorded variant specific *k*_cat_^app^ data, converting our 2D plausibility map into a 3D quantitative fitness landscape (**Figure 2B** and **Figure S11**). The resulting 3D view of the traversed fitness landscape captured the feature of wildtype resting in a fitness valley surrounded by multiple accessible fitness peaks, which were simultaneously explored during the directed evolution campaign. The best variant of the campaign (S68P-T241A-P203Q-H244Y) was identified as one of the few very high peaks of the landscape (**Figure 1F** and **Figure 2B**).

To reconstruct how the directed evolution campaign moved through sequence space, we ‘replayed the tape’ (*37*) of our time-resolved sequence-function data (**Figure 2C**). We found that evolution followed an explore-and-fix pattern with stepwise exploration of sequence space followed by fixation of successful variants. The first round explored possible single point mutations, forming a halo around the wildtype sequence (**Figure 2C panel 1**). To identify which single point variants dominated subsequent rounds, we extracted those with the highest node degree in the network graph, i.e., those with the highest number of direct descendants (**Figure 2C panel 2** and **Figure 2D**) (*35*). The active-site-adjacent mutation T241A clearly stood out, with a normalised node degree of 31 compared with 4 for the runner-up (**Figure 2D**). Analysis of the population’s mutational profile across rounds supported its subsequent dominance, with T241A occurring in almost 90% of round 5 variants (**Figure 2E** and **Figure S12**). Thus, we defined T241A as the first ‘founder’ variant of the evolution campaign: a variant from which a large number of viable one amino-acid substitution descendants were explored, thereby forming a well-connected hub for subsequent evolution in the network.

Accordingly, we identified the double mutant S68P-T241A, with a node degree of 30, as the second founder variant of the campaign (the runner-up node degree was 9, **Figure 2C panel 3** and **Figure 2D**). The founder role of S68P-T241A was also reflected at the population level, with almost 40% of all variants in round 5 containing the mutation S68P (**Figure 2E** and **Figure S12**). Finally, the two variants S68P-P203Q-T241A and S68P-T241A-H244Y were classified as potential third founders (**Figure 2C panel 4**). Altogether, the four observed founder variants formed a direct path from wildtype to the campaign’s fitness peak S68P-P203Q-T241A-H244Y (**Figure 2B** and **Figure S13**).

### Positive epistasis creates founders that rewire adaptive trajectories

We set out to identify the competitive advantage of founder variants and determine whether it arose from epistatic interactions that shaped subsequent adaptive trajectories. Overall, the dETM dataset allowed epistasis to be quantified for more than 461 interactions between 247 mutations (**Figure S14**). Among all double mutants explored from wildtype, 69% (109 of 158) could be explained by an additive model, with their measured fitness falling within 2-fold of the expected fitness based on their constituent single mutations (**Figure 3A**). This high combinability suggests that early evolution was not constrained by pervasive negative epistasis and traversed a generally smooth adaptive landscape. The remaining 31% of interactions deviated from additivity, including 39 (80%) positively epistatic double mutants, 14 of which contained T241A (blue dots in **Figure 3A**). Notably, the founder S68P-T241A showed one of the strongest interactions, with fitness 3.1-fold above the additive expectation (pink dot in **Figure 3A**). This positive epistatic interaction produced a 23-fold improvement in *k*_cat_ (observed *k*_cat_ of 106 s^-1^ vs expected *k*_cat_ of 35.4 s^-1^, **Figure 3E**, **Table S2** and **Figure S9**).

**Figure 3:**
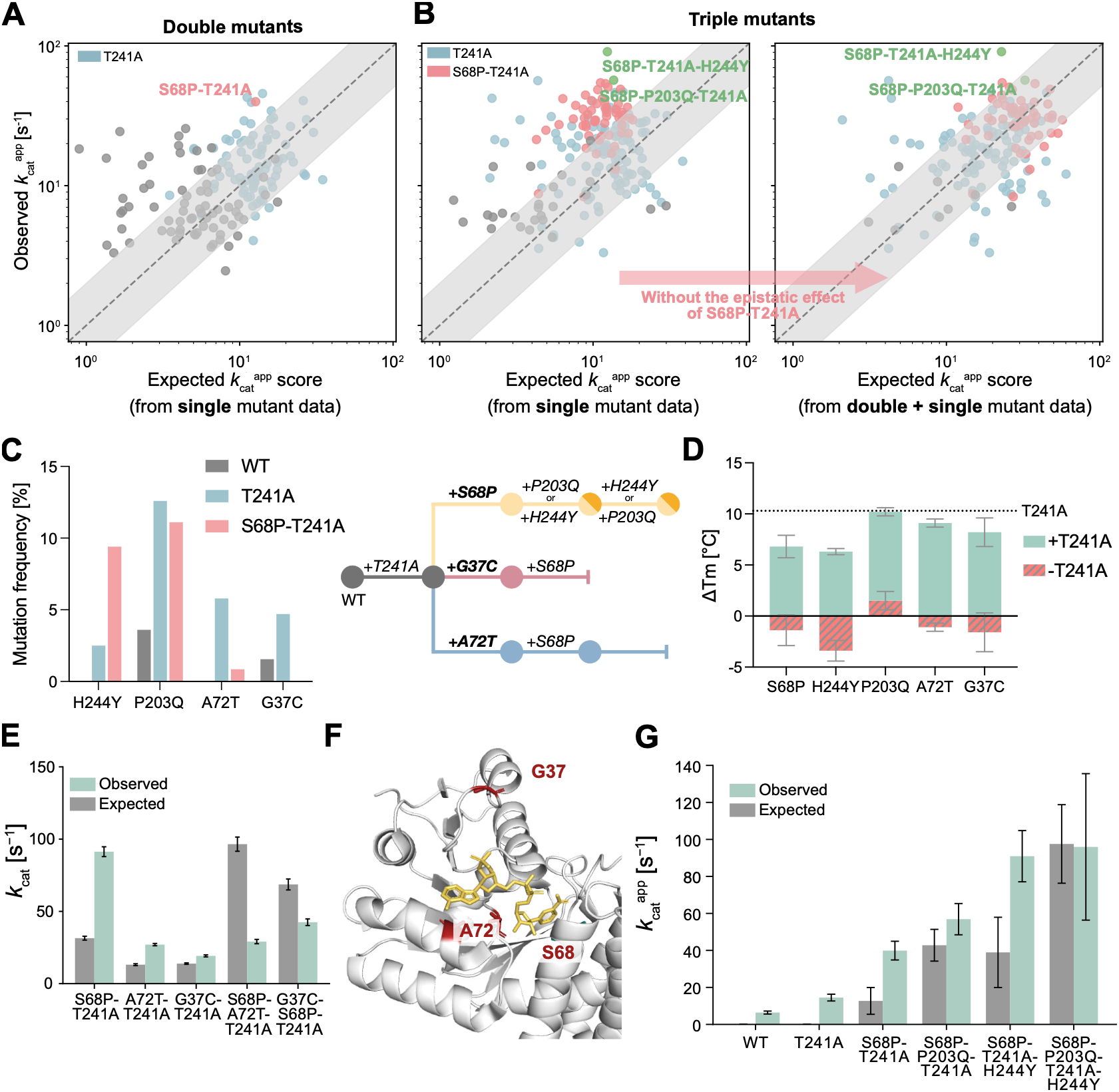
Epistasis shapes the directed evolution of *Sr*IRED. **(A)** Analysis of epistasis in double mutants. Most double mutants can be explained through additivity (± 2-fold of the additive expectation) with expected and observed fitness matching. 39 variants are positively epistatic among which 14 variants contain the mutation T241A (blue dots). One of the most fit and most positively epistatic mutants is S68P-T241A (pink dot). **(B)** Analysis of epistasis in triple mutants. A high proportion of highly active triple mutants is positively epistatic (left panel). Most of them contain the mutations S68P-T241A (pink dots). Excluding the single positive epistasis event between S68P and T241A, additivity can be restored (right panel). **(C)** Co-evolution of a selected set of mutations with the founder mutants T241A and S68P-T241A. Within the dETM dataset, the mutation frequency of the selected position was counted in the background of wildtype and the two founders (left). While all four analysed mutations enriched in the presence of T241A, only H244Y enriched further with S68P-T241A while P203Q stayed constant, and A72T and G37C de-enriched. This points to alternative trajectories being explored simultaneously (right). **(D)** The marginal detriment to stability of candidate single mutations is globally buffered by T241A. Melting temperatures T_M_ of mutations which co-evolved with T241A measured in thermal shift assays in three replicates and expressed as change in melting temperature compared to wildtype (ΔT_M,_ with T_M_^WT^ = 38.5 ± 0.6 °C). Errors represent standard deviations. **(E)** Epistatic relationship of T241A and S68P-T241A with dead-end alternative mutations. Michaelis-Menten kinetics were conducted with variable concentrations of cyclohexanone (0.1 - 52.7 mM), constant concentration of cyclopropylamine (75 mM) and NAPDH (1.5 mM), and variable enzyme concentrations (from 0.0015 to 0.05 g/L) in 100 mM Tris-HCl buffer at pH 8 and T = 25°C. Initial rates were calculated based on NADPH depletion at 380 nm using a standard curve of NADPH at that wavelength. **(F)** The Rossman fold of *Sr*IRED. The positions G37, S68, and A72 (marked in red) are located at the NADPH cofactor (marked in yellow) binding site and likely rivalling to interact with the cofactor (PDB code 5OCM^31^). **(G)** Positive epistasis of the founder variants. Larger observed fitness values compared to additive behaviour calculated from individual fitness scores suggests positive epistasis at each step towards the campaign’s fitness peak.

In the next evolutionary step, positive epistasis appeared to become more prevalent, increasing from 25% (39 of 158) of double mutants to 42% (79 of 186) of triple mutants (**Figure 3B panel 1)**. However, 53% of positively epistatic triple mutants contained the strongly epistatic S68P-T241A pair (pink dots in **Figure 3B panel 1)**, suggesting that the apparent increase in positive epistasis might largely reflect a single earlier founder event. When S68P-T241A was therefore treated as a single background, only 5% (3 of 57) of its triple-mutant descendants remained positively epistatic (pink dots in **Figure 3B panel 2**). Across the landscape, this reduced the fraction classified as positively epistatic from 42% to 12% (20 of 163). Thus, we show that a single founder event can propagate through subsequent genotypes, reshaping the epistatic topography of the fitness landscape along an adaptive evolutionary trajectory. Similar effects were observed for the two founders S68P-P203Q-T241A and S68P-T241A-H244Y and their influence on quadruple variant epistasis (**Figure S15**).

Next, we considered alternative trajectories leading from the first founder T241A and investigated mechanistically why they did not prevail in the campaign. Apart from the founder mutations S68P, P203Q and H244, two other mutations were significantly enriched in the background of T241A: A72T (5.8%) and G37C (4.7%), constituting two alternative trajectories within the campaign (**Figure 3C)**. However, in contrast to the other founders, these two mutations were then de-enriched in the prominent background of S68P-T241A (**Figure 3C**), indicating that they provide alternatives that are incompatible with the winning trajectory. Activity-improving mutations often have destabilising effects (*38–40*); therefore, we explored whether particular trajectories suffered from a stability collapse as a reason for their deselection. However, S68P, P203Q, H244Y, A72T, and G37C all had only marginally negative effects on stability on top of the 10.3 ± 0.5 °C increase in melting temperature conferred by T241A acting as a global suppressor (*41*) (**Figure 3D** and **Table S2**). Therefore, stability collapse does not explain the selection of the founder trajectory over its alternatives.

To test whether catalytic interactions instead explained founder selection, we measured Michaelis-Menten kinetics of the founder and dead-end mutants on T241A and S68P-T241A backgrounds. Both alternative mutantions displayed positive epistasis with T241A but negative epistasis with S68P-T241A (**Figure 3E)**. A structural model suggests that this switch arises from mutually exclusive interactions with the NADPH cofactor (**Figure 3F**). By contrast, the founder mutation H224Y showed strong positive epistasis with S68P-T241A, while P203Q remained additive (**Figure 3G**). These findings indicate that S68P-T241A rewired the enzyme’s epistatic network of the catalytic-rate landscape, mediated by the substrate recognition interface, such that the effects of previously beneficial mutations differed at the next evolutionary step (*42*). Positive epistasis therefore not only generated founder variants but redirected subsequent evolution: founder-compatible mutations persisted, whereas previously beneficial but incompatible mutations were out-competed as evolutionary dead-ends.

### Evolutionary past does not predict experimental future

Having reconstructed the campaign’s adaptive landscape and identified the epistatic interactions that shaped it, we asked whether the trajectories leading to the campaign’s hits could have been predicted from evolutionary sequence information. We therefore analysed natural homologues of *Sr*IRED. Protein-family multiple sequence alignments (MSAs) have previously been used to predict protein sequence plausibility and propose function-improving mutations by taking lessons from a protein’s ‘evolutionary past’ (*24, 38, 43, 44*). We retrieved the 5,000 closest homologues of *Sr*IRED using a BLAST search and constructed an MSA and a phylogenetic tree (**Figure S16**). In the laboratory evolution campaign, T241A emerged as the first founder variant and consequently alanine was the most common mutation at position 241 in the dETM dataset (**Figure 4A**). The MSA likewise identified alanine as the most frequent residue at position 241, showing that the first step in the laboratory campaign aligned with the natural sequence record (**Figure 4B**).

**Figure 4:**
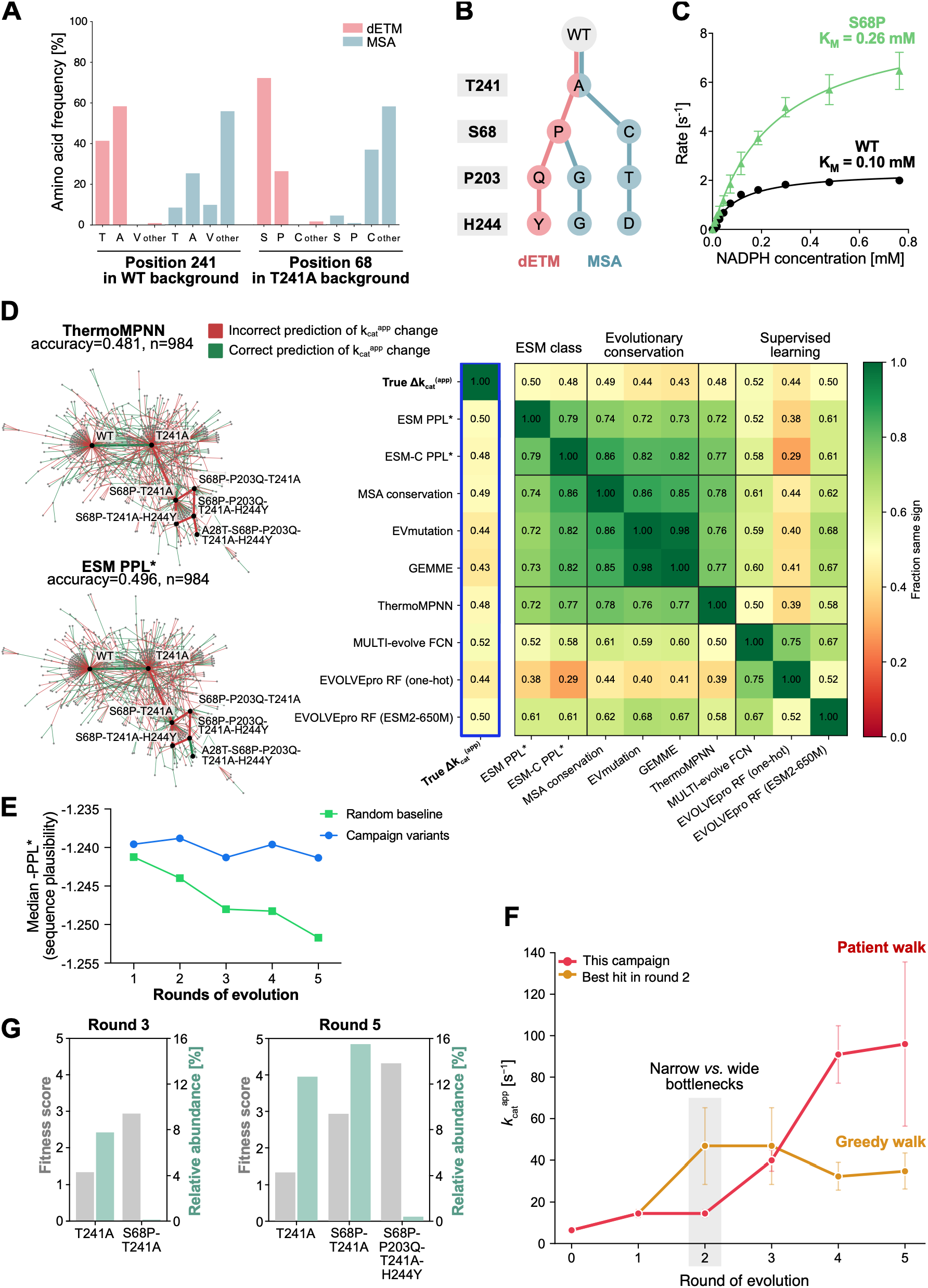
Experimental design influences the success of a directed evolution campaign. **(A)** Comparison of mutational profiles in the dETM dataset and an MSA of 5000 *Sr*IRED homologues retrieved from a BLAST search. The position 241 was analysed in the dETM and MSA datasets and the most frequent amino acids are displayed (left). Both datasets were filtered for sequences with an alanine at position 241 and respective subsets were used for analysis of the most frequent amino acids at position 68 (right). **(B)** Phylogenetic tree constructed from the dETM and MSA data. The order in which the four positions 241, 68, 203, and 244 were mutated was chosen based on the directed evolution campaign data. For each position, the most frequent mutation was chosen from the dETM and MSA dataset to construct a phylogeny. Two possible trajectories emerged for the MSA data and one single possible trajectory emerged for the laboratory evolution campaign. **(C)** Michaelis-Menten kinetics of *Sr*IRED wildtype and the mutant S68P with variable concentrations of NADPH and constant concentrations of cyclohexanone (30 mM) and cyclopropylamine (30 mM) in 100 mM Tris-HCl buffer (pH 8). Initial rates were calculated based on NADPH depletion at 340 nm and T = 25°C. Kinetics revealed a 2.6-fold increased K_M_ from 0.10 ± 0.01 mM for wildtype to 0.26 ± 0.03 mM for the mutant. **(D)** Models’ ability to predict the direction of *k* ^app^ change between variants connected in the network graph of the campaign (left). The accuracy (proportion of correct predictions) is similar between models but does not exceed the random baseline of 0.5 (right). **(E)** Variant plausibility estimation using ESM2. Variants from each round were assessed for their plausibility using the negative median model perplexity -PPL (blue). For every observed variant, five random genotypes were sampled from a round-specific substitution pool with the same mutational load as in the observed genotype. These were passed through the model to provide a random baseline scenario (green). **(F)** Performance of different selection regimes over multiple rounds of directed evolution. Using the recorded dETM data, two selection regimes were simulated by applying a stringent and a permissive selection threshold to the population in round 2. The stringent selection threshold selected the best variant of round 2 and was representative of traditional directed evolution campaigns which rely on *greedy walks*. The permissive selection threshold followed the winning trajectory of the campaign presented in this study where a pool of variants was carried forward each round. In round 2, a variant which was ranked 31^st^ in fitness was selected for the winning trajectory, representing a *patient walk* in sequence space. **(G)** Effect of rescreening depth on hit identification. Fitness score and relative abundance within the full variant pool were compared for T241A, S68P-T241A, and S68P-P203Q-T241A-H244Y in round 3 (left panel) and round 5 (right panel). Relative abundance, which determines the likelihood of detecting a variant during rescreening, was calculated as the number of NGS counts assigned to that variant divided by the total number of counts in the respective round.

We then focused on the next step of the campaign, which introduced a mutation of serine at position 68. In the dETM dataset, proline was the most common substitution at this position, whereas the MSA showed cysteine as the most frequent residue (occurring in 40% of the analysed IREDs, **Figure 4A**). In the campaign, S68P occurred often in the background of T241A, due to a strong positive epistatic interaction. However, this combination was rare in the MSA: only 0.7% of sequences contained P68-A241, compared to 2.2% containing P68-T241 (**Figure 4A** and **Figure S20**). The laboratory trajectory therefore diverged from the natural sequence record at S68P (**Figure 4B**). Subsequent laboratory steps introduced P203Q and H244Y. Natural homologues instead favoured threonine or glycine at position 203 and aspartate or glycine at position 244 (**Figure 4B**).

This bifurcation between laboratory and natural evolution may reflect selection for different enzymological traits (*45*): the laboratory campaign selected primarily for catalytic rate under substrate-saturating conditions (**Figure S7**), whereas natural evolution might have been driven by catalytic efficiency at low substrate concentrations. S68P exemplified this trade-off. Its positive epistatic interaction with T241A generated a strong catalytic-rate advantage (**Figure 3E**), but S68P also induced a 2.6-fold increase in K_M_ for NADPH, from 0.10 mM in wildtype to 0.26 mM in the mutant (**Figure 4C** and **Figure S19**). During laboratory evolution, a saturating NADPH concentration of 5 mM was used, masking this cofactor-affinity cost.

Similarly, S68P induced a trade-off between *k*_cat_ and K_M_ for cyclohexanone (**Figure S20**). Together, these observations suggest that, under saturating laboratory conditions, the catalytic-rate advantage of S68P-T241A outweighed its affinity costs. This enabled the founder lineage to expand and outcompete alternative lineages. By contrast, the lower cofactor and substrate concentrations likely encountered during natural evolution (*46*) would expose these affinity costs and impose counter-selection pressure against S68P. This provides a plausible explanation for the rarity of P68-A241 in the MSA.

However, simple inspection of a MSA may miss residue interdependencies, co-evolution, sequence-structure constraints and enzyme-specific contingencies. We therefore asked whether machine learning models drawing on broader evolutionary, structural, or experimental information could have predicted the campaign’s outcome. We benchmarked computational approaches spanning evolutionary sequence models (ESM, EVmutation and GEMME) (*23, 24, 47*), structure-based stability prediction (ThermoMPNN) (*48*), and supervised learning from experimental data (EVOLVEpro and MULTI-evolve) (*25, 26*) against the individual mutational steps of the laboratory evolution campaign captured in the dETM dataset.

Using a network graph of the campaign, we labelled each edge between two variants according to whether the *k* ^app^ increased or decreased during evolution (**Figure 4D** and **Figure S20**). We then asked whether each model could predict the direction of change and identify productive evolutionary steps. None of the tested methods exceeded the random baseline scenario (**Figure 4D**). Thus, we concluded that the winning trajectory of the directed evolution campaign could not have been predicted through computational methods. We furthermore observed a strong agreement among several models, despite their inability to predict improvements, highlighting the risk associated with *in silico* model evaluation using cross-model performance, instead of benchmarking models on experimental datasets (*49, 50*). This stands in stark contrast to the success of cross-model filtering in the design of *de novo* binding proteins (*51*).

Since the models did not reliably predict productive evolutionary steps, we asked whether unsupervised protein language models might have learnt a holey landscape in which variants are assigned binary activity labels: active or inactive (*52*). We compared ESM zero-shot predictions for the variants observed across five rounds of evolution with those for random genotypes matched to each round’s average variant Hamming distance. Random genotypes became progressively less plausible with increasing Hamming distance, whereas the observed dETM variants retained similar plausibility across rounds (**Figure 4E**). This pattern suggests that protein language models like ESM act as plausibility filters for enzymes, rather than predictors of catalytic fitness.

### Permissive selection and high-throughput screening enable patient walks to fitness peaks

We next asked whether the dETM dataset could be used to further illuminate the black box of directed evolution and deduce experimental design rules for future campaigns. To this end, we assessed how selection thresholds and screening depth shaped the outcome of the recorded campaign. In our evolutionary system, microfluidic screening evaluated the catalytic rate of each variant as *k* ^app^ and selected the best 200-500 variants from libraries comprising approximately 200,000 variants per round (**Figure 1C**). We applied an initially permissive selection regime that became progressively more stringent, with the effective *k* ^app^ selection threshold increasing from 0.6-fold to 1.2-fold relative to wildtype (**Figure S21**). After each round, the entire selected pool was diversified with one or two additional amino acid substitutions (**Figure 1A)**. Conceptually, this permissive selection regime resembles a beam search, preserving multiple genetic backgrounds and allowing parallel evolutionary trajectories to be explored (**Figure 2A**).

To place this design in context, we retrospectively replayed the campaign using the dETM records under our permissive regime and single-parent greedy search in which only the best performing variant is re-diversified (*53, 54*). These alternatives have frequently produced diminishing returns in multi-round campaigns (*9, 55, 56*). Our permissive selection regime recovered the winning trajectory to the fitness peak (red line in **Figure 4F**). The path included T241A, which was retained in round 2 despite ranking only 31^st^ by fitness (**Table S3**). Only due to this permissive selection event, S68P-T241A could emerge in round 3 and initiated the winning trajectory (**Table S3**). By contrast, the single-parent greedy replay retained only the best performing variant in each round and restricted the next step to its recorded descendants. This selected K38N-E210G-T241A in round 2, which had few descendants showing only marginal further improvements in *k* ^app^ (blue line in **Figure 4F** and **Table S3**). Thus, the two selection regimes diverged in round 2: fixing the rounds winner K38N-E210G-T241A led to diminishing returns following its progeny, whereas retaining the 31^st^-ranked T241A opened the path to the campaign’s fitness peak. We therefore described our selection regime as a *patient walk* through sequence space: a strategy which retains backgrounds with moderate improvements and unlocks access to trajectories with greater potential for future adaptation.

The ability of a patient walk to retain multiple lineages comes at the cost of an increased screening burden. We therefore asked whether high-throughput screening not only supported our permissive selection regime but also increased the probability of recovering rare hits. To test this, we used NGS counts to quantify the relative abundance of each variant in the selected pools and estimate its probability of recovery during clonal rescreening. In round 3, T241A was the most abundant variant, accounting for 7.7% of the pool whereas S68P-T241A showed a ninefold improvement in *k*_cat_^app^ over T241A but accounted for only 0.1% of the NGS counts (**Figure 4G** and **Figure S21**). A similar pattern was observed in round 5: S68P-P203Q-T241A-H244Y had the highest fitness score of 4.3 but accounted for only 0.1% of the variant pool, whereas the less fit variants T241A and S68P-T241A accounted for 12.6% and 15.4% respectively (**Figure 4G** and **Figure S22**). This discrepancy between fitness and genotype abundance reflects a temporal lag in enrichment: newly emerging hits can remain rare for several rounds, whereas previously selected, lower-fitness variants remain abundant. Consequently, recovery during low-throughput clonal rescreening is governed by genotype frequency rather than fitness. At the observed round 5 frequencies, oversampling statistics indicate that approximately 23 clones would be sufficient to recover either T241A or S68P-T241A with at least 95% probability when considered individually (*57*). In contrast, recovering S68P-P203Q-T241A-H244Y would require screening approximately 2,600 clones, equivalent to 27 96-well plates (**Table S4**). Screening more than 10^5^ variants by microfluidics increased the probability of capturing such rare hits (*58*) and deep sequencing enabled their identification without exhaustive clonal re-screening.

## Implications and Conclusions

The dETM framework reveals an evolutionary solution to the numbers problem in the exploration of function in sequence space: how to deal with the combinatorial explosion of possible variants that would never be experimentally screenable. Here, we observe ‘founders’ that emerge as focal points and highly connected hubs able to traverse sequence space based on combinable mutational patterns. These provide directionality and connectivity towards higher activity through positively epistatic interactions and make an onward trajectory more likely by outcompeting ultimately frustrated pathways without onward connection. We note an analogy to another of Nature’s marginal chance problems, protein folding. In the case of folding, partially-folded intermediates and ensembles have been invoked that ‘funnel’ folding pathways towards the marginal chance outcome of a native structure, allowing folded proteins to be a prevalent feature in living systems (*59*). Likewise founders funnel mutants towards constructive trajectories, thus beating the odds stacked against identifying the few functional proteins.

DMS was repurposed in our campaign to provide insight into the dynamics of the exploration process itself, rather than its usual application of locally exploring the fitness landscape immediately surrounding a wild-type protein (i.e. single amino acid mutations) (*14, 36, 60*). The fitness-labelled data obtained from multiple rounds of screening illuminate the currently often obscure black box process of directed evolution when moving several mutations away from the starting point. Such insight will provide the basis for troubleshooting, when evolution is unsuccessful or limited by diminishing returns scenarios (*61*). Next generation sequencing analysis of the evolving population allowed us to identify not just single hits, but larger numbers of mutants and subpopulations. Even if these are not yet dominant, their round-to-round enrichments highlight successful dynamic adaptation. While the most dynamic trajectories may become dominant later, dETM identifies these lagging upstarts early. Ultrahigh throughput and deep sequencing are necessary to reveal deep and dynamic information, where low throughput plate screening is only able to give a shallow and static picture. At the same time, the challenges caused by the abundance of wild type in sparsely mutated (*36, 60*) libraries or, more generally, skewed library distributions (*62*) that plague conventional directed evolution campaigns can be analysed and evaluated. The greater transparency of dETM provides novel insight into the areas of sequence space characterised by higher or lower evolvability potential.

Conceptually different directed evolution strategies can now also be evaluated. By using a freely adjustable selection threshold in a screening assay we can control the size of the selected sub-population to deliberately carry a larger or smaller sequence footprint forward from round to round. Tweaking the stringency of rounds is an experimental variable that can now be rationalised (and optimised) based on sequencing readouts, suggesting that DMS information in dETM can spawn build-test-learn cycles. Likewise, comparing and contrasting different mutation rates or library synthesis methods will lead to evaluation of the success of strategies based on data rather than following *a priori* hypotheses that can now be tested instead.

The models of global diminishing returns and epistasis predict that beneficial mutations should, on average, confer progressively smaller effects on fitter backgrounds (*63–65*). In contrast, our IRED trajectory shows sustained improvement without a systematic decline in effect size: individual mutations gain or lose value depending on their genetic background, while neither catalytic activity nor the combined activity–stability phenotype approaches an obvious evolutionary ceiling, in contrast to the strong diminishing returns previously observed during laboratory enzyme evolution (*9*). These observations suggest that patient walks, enabled through permissive selection and high-throughput screening, de-risk landscape traversal by simultaneously exploring multiple evolutionary trajectories. They thereby leverage beneficial interactions that help negotiating shifts in evolvability that would otherwise become entrenched in traditional greedy search strategies. As a result, directed evolution is constrained less by a universal diminishing-returns regime than by shifting epistatic networks, consistent with the broader observation that the effects of individual mutations can depend strongly on genetic background rather than simply scaling with starting fitness (*65*). Such epistatic interactions, in this example present at every step and propelling the campaign forward, may present the ultimate barrier to the computational prediction of evolutionary outcomes, as epistatic drift (*66*) idiosyncratically modulates mutational effects in long-term sequence space traversals.

Using dETM, we find that existing computational methods are poor predictors of campaign trajectories, instead acting merely as ‘plausibility filters’ that allow discrimination between random mutants and productive campaign members, yet without the ability to navigate towards higher activity: this challenge is readily met by experimental evolution in the dETM format. These observations resolve the paradox of how computational strategies trained on evolutionary data are often effective at protein design and re-design (*47*), but generally unable to guide designs or discovery campaigns reliably to fitness maxima without additional experimental input (*25, 67*). 2D sequence coordinates of a fitness landscape (i.e. the x and y axes of a 3D space, **Fig. 2B**) are predictable with respect to function retention, but its third dimension (the activity axis) can only be elaborated by experimental data of the type that we provide with dETM: laboratory evolution reveals computationally unpredictable epistatic trajectories with functional quantification as ground truth for engineering or computational re-evaluation.

The analytical framework presented here holds the potential to yield understanding and ultimately gain control of trajectories through sequence space towards functional improvements by providing datasets ready for AI/ML interpretation or for classical rational mechanistic analysis. While we describe *one* scenario involving founders as a key feature, other proteins may evolve by different mechanisms – if Nature is indeed a tinkerer (*68*), it is likely that a number of mechanisms will emerge. Currently, practical insights into improving the evolvability of a protein are scant (*69*). A stable starting point is useful (*70*), and accommodation of mutational load has been highlighted as basis of the ‘*one amino acid at a time*’ strategy (*36, 60*), but dETM highlights and characterises the evolving populations that are sufficient for a constructive trajectory. dETM provides insights on populations and their dynamics, not just sequences of winners. In the future, dETM can be used as an experimental framework to quantify evolvability of proteins by evaluating the viability of dynamic walks in sequence space (*69, 71–74*), with properties such as starting point thermostability (*22, 60, 70, 75*), ancestral reconstruction (*76–78*), selection threshold stringency (*79*), library generation concepts (*25, 36, 80–82*) and protein design strategies (*70, 83*) all becoming quantifiable in their effects on a fitness-mapped evolutionary network. Likewise, the intrinsic properties that make a protein more or less evolvable such as stability, conformational dynamics, frequency and nature of epistatic interactions (creating rugged or smooth fitness landscapes) can be probed. We know that these parameters are important, but now we offer a readout that can report on how proteins respond to challenges under different parametric settings. In the future the ability to record and evaluate comprehensive trajectories will enable quantification of the roles of hitherto poorly constrained choices that make or break protein engineering attempts. Datasets recorded using this methodology, combined with deliberately altering environmental or protein-intrinsic parameters, will provide case studies of frustrated or successful trajectories that can shed light on ground truths of protein engineering left so far in the dark and reveal real fitness landscapes as long anticipated by John Maynard Smith (*84*). Comparative studies based on dETM will be able to use a common language for inter-campaign comparison, both via network statistics for identification of hub or dead-end mutants, and benchmarking statistics of predictive tools for protein evolution to fulfil the hitherto elusive task of reliably reaching high functional enzymatic efficiency.

## Acknowledgements

We thank the members of the Hollfelder group for helpful discussions and sequencing facility at the University of Cambridge Department of Biochemistry for their support.

## Author Contributions

Conceptualisation: FEHN, ML, MP, MG, FH

Methodology: FEHN, MP, MS, MG, EJM, FH

Formal analysis: FEHN, ML, MP, MS, SVM, SC

Investigation: FEHN, ML, MP, MS, HB, CS, SVM, SC

Data Curation: FEHN, ML, MS

Writing – Original Draft: FEHN

Writing – Review, Editing & Rewriting: FEHN, ML, MP, FH

Visualisation: FEHN, ML, MP, MS

Funding acquisition: FH

## Data, code, and materials availability

Data processing scripts are stored on a GitHub repository and deposited at Zenodo (10.5281/zenodo.23805430). Raw sequencing files were uploaded to ENA (Study accession: PRJEB129111).

