## Supplementary Information for "Deep sequence-function mapping reveals computationally unpredictable epistatic trajectories that drive laboratory evolution"

### Materials and Methods

#### A. Molecular biology

<u>Cloning of *Sr*IRED wildtype</u>

The gene encoding for *Sr*IRED (a bacterial IRED from *Streptosporangium* roseum) was cloned into a pRSF vector with N-terminal $His_6$-tag, and under control of a T7 promoter.

##### *SrIRED nucleotide sequence:*

ATGGGCAGCAGCCATCATCATCATCATCACAGCAGCGGCCTGGTGCCGCGCGGCAGCCATATGCGGGACACCGATGTCACTGTCCTCGGCCTGGGTCT
GATGGGCCAGGCTTTGGCAGGCGCCTTTCTCAAAGATGGGCATGCGACGACCGTTTGGAATCGTTCTGAAGGCAAAGCCGGTCAGCTGGCGGAACAG
GGTGCGGTACTGGCCAGCAGCGCCCGGGATGCGGCCGAGGCGAGCCCACTGGTAGTTGTTTGCGTTTCAGACCACGCCGCGGTTCGCGCGGTGCTG
GATCCTCTGGGTGATGTTCTGGCGGGGCGTGTTCTGGTGAACCTCACCTCTGGAACTAGCGAACAGGCCCGCGCCACGGCGGAGTGGGCCGCTGAAC
GCGGGATTACATACCTTGACGGAGCTATCATGGCAATCCCGCAGGTTGTCGGCACAGCAGATGCATTCTTACTGTACTCTGGGCCGGAAGCCGCTTATG
AGGCGCACGAACCGACCTTACGGAGCCTGGGGGCGGGTACCACTTATTTGGGCGCCGACCACGGCCTGTCATCTTTGTATGACGTCGCGTTGCTCGGC
ATCATGTGGGGCACGCTGAATTCTTTCCTCCACGGGGCGGCATTGTTGGGCACAGCCAAAGTGGAAGCGACGACCTTTGCACCGTTCGCGAACCGTTG
GATTGAAGCGGTAACCGGGTTTGTATCCGCGTATGCTGGTCAGGTGGACCAGGGGGCCTACCCGGCACTGGACGCGACTATCGATACCCACGTGGCTA
CGGTAGATCATCTGATTCACGAGTCCGAGGCCGCTGGCGTGAACACGGAGCTCCCACGCCTCGTGCGCACGCTGGCCGACCGTGCCCTGGCGGGAG
GCCAGGGCGGTCTTGGTTACGCCGCTATGATTGAGCAGTTTCGGTCTCCTAGCTAA

##### *SrIRED whole plasmid sequence:*

ctgtaacatcattggcaacgctacctttgccatgtttcagaaacaactctggcgcatcgggcttcccatacaatcgatagattgtcgcacctgattgcccgacattatcgcgagcccatttatacccatataaatcag
catccatgttggaatttaatcgcggcctagagcaagacgtttcccgttgaatatggctcatactcttcctttttcaatattattgaagcatttatcagggttattgtctcatgagcggatacatatttgaatgtatttagaaaa
ataaacaaataggcatgcagcgctcttccgcttcctcgctcactgactcgctacgctcggtcgttcgactgcggcgagcggtgtcagctcactcaaaagcggtaatacggttatccacagaatcaggggataaa
gccggaaagaacatgtgagcaaaaagcaaagcaccggaagaagccaacgccgcaggcgtttttccataggctccgcccccctgacgagcatcacaaaaatcgacgctcaagccagaggtggcgaaa
cccgacaggactataaagataccaggcgtttccccctggaagctccctcgtgcgctctcctgttccgaccctgccgcttaccggatacctgtccgcctttctcccttcgggaagcgtggcgctttctcatagctcacg
ctgttggtatctcagttcggtgtaggtcgttcgctccaagctgggctgtgtgcacgaaccccccgttcagcccgaccgctgcgccttatccggtaactatcgtcttgagtccaacccggtaagacacgacttatcgc
cactggcagcagccattggtaactgatttagaggactttgtcttgaagttatgcacctgttaaggctaaactgaaagaacagattttggtgagtgcggtcctccaacccacttaccttggttcaaagagttggtagct
cagcgaaccttgagaaaaccaccgttggtagcggtggtttttctttatttatgagatgatgaatcaatcggtctatcaagtcaacgaacagctattccgttactctagatttcagtgcaatttatctcttcaaatgtagca
cctgaagtcagccccatacgatataagttgtaattctcatgttagtcatgccccgcgcccaccggaaggagctgactgggttgaaggctctcaagggcatcggtcgagatcccggtgcctaatgagtgagctaa
cttacattaattgcgttgcgctcactgcccgctttccagtcgggaaacctgtcgtgccagctgcattaatgaatcggccaacgcgcggggagaggcggtttgcgtattgggcgccagggtggtttttcttttcaccagt
gagacgggcaacagctgattgcccttcaccgcctggccctgagagagttgcagcaagcggtccacgctggtttgccccagcaggcgaaaatcctgtttgatggtggttaacggcgggatataacatgagctgt
cttcggtatcgtcgtatcccactaccgagatgtccgcaccaacgcgcagcccggactcggtaatggcgcgcattgcgcccagcgccatctgatcgttggcaaccagcatcgcagtgggaacgatgccctcatt
cagcatttgcatggtttgttgaaaaccggacatggcactccagtcgccttcccgttccgctatcggctgaatttgattgcgagtgagatatttatgccagccagccagacgcagacgcgccgagacagaacttaat
gggcccgctaacagcgcgatttgctggtgacccaatgcgaccagatgctccacgcccagtcgcgtaccgtcttcatgggagaaaataatactgttgatgggtgtctggtcagagacatcaagaaataacgcc
ggaacattagtgcaggcagcttccacagcaatggcatcctggtcatccagcggatagttaatgatcagcccactgacgcgttgcgcgagaagattgtgcaccgccgctttacaggcttcgacgccgcttcgttct
accatcgacaccaccacgctggcacccagttgatcggcgcgagatttaatcgccgcgacaatttgcgacggcgcgtgcagggccagactggaggtggcaacgccaatcagcaacgactgtttgcccgcca
gttgttgtgccacgcggttgggaatgtaattcagctccgccatcgccgcttccactttttcccgcgttttcgcagaaacgtggctggcctggttcaccacgcgggaaacggtctgataagagacaccggcatactct
gcgacatcgtataacgttactggtttcacattcaccaccctgaattgactctcttccgggcgctatcatgccataccgcgaaaggttttgcgccattcgatggtgtccgggatctcgacgctctcccttatgcgactcct
gcattaggaaattaatacgactcactataggggaattgtgagcggataacaattcCCCTGTAGAAATAATTTTGTTTAACTTTAATAAGGAGATATACCATGGGCAGCAGC
CATCATCATCATCATCACAGCAGCGGCCTGGTGCCGCGCGGCAGCCATATGCGGGACACCGATGTCACTGTCCTCGGCCTGGGTCTGATGG
GCCAGGCTTTGGCAGGCGCCTTTCTCAAAGATGGGCATGCGACGACCGTTTGGAATCGTTCTGAAGGCAAAGCCGGTCAGCTGGCGGAACA
GGGTGCGGTACTGGCCAGCAGCGCCCGGGATGCGGCCGAGGCGAGCCCACTGGTAGTTGTTTGCGTTTCAGACCACGCCGCGGTTCGCG
CGGTGCTGGATCCTCTGGGTGATGTTCTGGCGGGGCGTGTTCTGGTGAACCTCACCTCTGGAACTAGCGAACAGGCCCGCGCCACGGCGG
AGTGGGCCGCTGAACGCGGGATTACATACCTTGACGGAGCTATCATGGCAATCCCGCAGGTTGTCGGCACAGCAGATGCATTCTTACTGTAC
TCTGGGCCGGAAGCCGCTTATGAGGCGCACGAACCGACCTTACGGAGCCTGGGGGCGGGTACCACTTATTTGGGCGCCGACCACGGCCTG
TCATCTTTGTATGACGTCGCGTTGCTCGGCATCATGTGGGGCACGCTGAATTCTTTCCTCCACGGGGCGGCATTGTTGGGCACAGCCAAAGT
GGAAGCGACGACCTTTGCACCGTTCGCGAACCGTTGGATTGAAGCGGTAACCGGGTTTGTATCCGCGTATGCTGGTCAGGTGGACCAGGGG
GCCTACCCGGCACTGGACGCGACTATCGATACCCACGTGGCTACGGTAGATCATCTGATTCACGAGTCCGAGGCCGCTGGCGTGAACACGG
AGCTCCCACGCCTCGTGCGCACGCTGGCCGACCGTGCCCTGGCGGGAGGCCAGGGCGGTCTTGGTTACGCCGCTATGATTGAGCAGTTTC
GGTCTCCTAGCTAATCTGGTAAAGAAACCGCTGCtgcgaaatttgaacgccagcacatggactcgtctactagcgcagcttaattaacctaggctgctgccaccgctgagcaataacta
gcataaccccttggggcctctaaacgggtcttgaggggttttttgctgaaacctcaggcatttgagaagcacacggtcacactgcttccggtagtcaataaaccggtaaaccagcaatagacataagcggctatt
taacgaccctgccctgaaccgacgacaagctgacgaccgggtctccgcaagtggcacttttcggggaaatgtgcgcggaacccctatttgtttatttttctaaatacattcaaatatgtatccgctcatgaattaattc
ttagaaaaactcatcgagcatcaaatgaaactgcaatttattcatatcaggattatcaataccatatttttgaaaaagccgtttctgtaatgaaggagaaaactcaccgaggcagttccataggatggcaagatcc
tggtatcggtctgcgattccgactcgtccaacatcaatacaacctattaatttcccctcgtcaaaaataaggttatcaagtgagaaatcaccatgagtgacgactgaatccggtgagaatggcaaaagtttatgca
tttctttccagacttgttcaacaggccagccattacgctcgtcatcaaaatcactcgcatcaaccaaaccgttattcattcgtgattgcgcctgagcgagacgaaatacgcggtcgctgttaaaaggacaattacaa
acaggaatcgaatgcaaccggcgcaggaacactgccagcgcatcaacaatattttcacctgaatcaggatattcttctaatacctggaatgctgttttcccggggatcgcagtggtgagtaaccatgcatcatca
ggagtacggataaaatgcttgatggtcggaagaggcataaattccgtcagccagtttagtctgaccatctcat

##### *SrIRED amino acid sequence:*

MGSSHHHHHHSSGLVPRGSHMRDTDVTVLGLGLMGQALAGAFLKDGHATTVWNRSEGKAGQLAEQGAVLASSARDAAEASPLVVVCVSDHAAV
RAVLDPLGDVLAGRVLVNLTSGTSEQARATAEWAAERGITYLDGAIMAIPQVVGTADAFLLYSGPEAAYEAHEPTLRSLGAGTTYLGADHGLSSLYD
VALLGIMWGTLNSFLHGAALLGTAKVEATTFAPFANRWIEAVTGFVSAYAGQVDQGAYPALDATIDTHVATVDHLIHESEAAGVNTELPRLVRTLAD
RALAGGQGGLGYAAMIEQFRSPS*

##### *SrIRED protein sequence without tag:*

MRDTDVTVLGLGLMGQALAGAFLKDGHATTVWNRSEGKAGQLAEQGAVLASSARDAAEASPLVVVCVSDHAAVRAVLDPLGDVLAGRVLVNLTS
GTSEQARATAEWAAERGITYLDGAIMAIPQVVGTADAFLLYSGPEAAYEAHEPTLRSLGAGTTYLGADHGLSSLYDVALLGIMWGTLNSFLHGAALL
GTAKVEATTFAPFANRWIEAVTGFVSAYAGQVDQGAYPALDATIDTHVATVDHLIHESEAAGVNTELPRLVRTLADRALAGGQGGLGYAAMIEQFRS
PS*

Cloning of *Sr*IRED variants

*Sr*IRED variants were cloned using site-directed ligation-independent mutagenesis (SLIM)(*1*), Gibson assembly(*2*), or using dial out PCR from a pool of plasmids or a plasmid library with primers specific to the UMI region(*3*). Cloning success was confirmed by Sanger sequencing.

Error-prone PCR library cloning of *Sr*IRED from wildtype

The epPCR library of *Sr*IRED wildtype was generated as described previously(*4*). The *Sr*IRED gene was amplified by error-prone PCR (epPCR) using the GeneMorph II random mutagenesis kit (Agilent) according to the manufacturer's protocol, with 2.4 µg of template plasmid and 25 PCR cycles. The resulting product was digested overnight with DpnI (NEB), extracted from an agarose gel, and further purified using SPRIselect beads (Beckman Coulter) following the manufacturer's instructions. A unique molecular identifier (UMI) was incorporated from randomized oligonucleotides (IDT) via three cycles of PCR using a primer annealing to the 3'end to generate double-stranded DNA. For library assembly, the following PCR fragments were generated: (i) promoter region amplified with primers GA1 and GA5, (ii) vector backbone amplified with primers GA2 and GA6, and (iii) the *Sr*IRED gene fragment amplified with primers GA3 and GA4 under error-prone conditions. Fragments were assembled using Gibson assembly mix, added on ice, and incubated at 50 °C for 1 h. The assembled product was purified with SPRIselect beads and transformed into electrocompetent E. coli (Lucigen). Transformation yielded a plate with approximately 60,000 colonies by dilution series.

*Primers for Gibson assembly of SrIRED epPCR library:*

GA1: atcccgcgaaattaatacgactcactatatgcgactcctgcattaggaaattaatacGACTCACTATAGGGG

GA2: agatctcgatcctctacgccggacgcatctaagggagagcgtcgagatcc

GA3: CCGCGCGGCAGCCATATG

GA4: caaatttcgcaGCAGCGGTTTCTTTACCAGATTA

GA5: CATATGGCTGCCGCGCGGCACCAG

GA6: TAATCTGGTAAAGAAACCGCTGCtgc

*UMI sequence:*

gatgcgtccggcgtagaggatcgagatctgNNNNNNNNNNGATCNNNNNNNNNNGATCNNNNNNNN NNGATCNNNNNNNNNNGATCNNNNNNNNNNgatcccgcgaaattaatacgactcactata

*Primer for UMI:*

tatagtgagtcgtattaatttcgcgg

Error-prone PCR library cloning of *Sr*IRED during directed evolution

The *Sr*IRED epPCR libraries produced during the directed evolution campaign were prepared by Golden-Gate assembly of the following four parts: (1) the *Sr*IRED gene amplified with error-prone PCR (epPCR) from the entire sorted variant pool of the respective previous round, (2) the unique-molecular-identifier (UMI) which provides a unique barcode to each variant in the library, (3) the T7 promoter, and (4) the pRSF backbone including a barcode (BC) which is specific for each round and each substrate. The error-prone PCR was performed using the GeneMorph II random mutagenesis kit (Agilent), following the instructions of the manufacturer. Conditions were 25 PCR cycles and 2.4 µg template plasmid (GGA5 & GGA6). The epPCR product was DpnI (NEB) digested overnight and purified using gel-electrophoresis. The UMI was ordered as single stranded oligonucleotide (IDT). For generation of the double stand, a primer was annealed at the 3' end under hybridization conditions. Subsequently, extension was performed using the large (Klenow) fragment of DNA Polymerase I (NEB). The final product was purified using SPRIselect beads (Beckman coulter) following the manufacturer's instructions. Vector (GGA1 & GGA2) and promoter region (GGA3 & GGA4) were produced by PCR. Fragments were assembled using golden gate assembly mix using the manufacturer's instructions (NEB) using SapI enzyme. After Golden-Gate assembly, the product was purified using SPRIselect beads and transformed into electrocompetent *E. coli* (E.cloni 10G Elite, Cambridge Bioscience). Quality of the produced library was confirmed by Sanger sequencing of ten random transformants.

*Primers for Golden-Gate assembly of SrIRED epPCR library:*
GGA1: ctatGCTCTTCattgaacgccagcacatggac
GGA2: cgttGCTCTTCtagctGCTCGGAGATTGAGAAAGCAgcatctaagggagagcgtcgag
GGA3: attgGCTCTTCgatatgcgactcctgcattaggaaat
GGA4: gataGCTCTTCaGGAACCAGGCCGCTGCTG
GGA5: ctagGCTCTTCaTCCGCGCGGCAGCCATATG
GGA6: cggtGCTCTTCacaaatttcgcaGCAGCGGTTTCTTTACCAGATTA

*UMI sequence:*
cactGCTCTTCagctgNNNNRYNNNNGATCNNNNRYNNNNGATCNNNNRYNNNNGATCNN
NNRYNNNNGATCNNNNRYNNNNgatcacgcgtcatgcatgcgactcactataaGAAGAGCacag

*Primer for UMI:*
ctgtGCTCTTCttatagtgagtcgcatgcatgacgcgtgatc

*Barcode (BC) for round 1:*
CCGGCGTAGAGGATCGAGAT

*Barcode (BC) for round 2:*
AAGTAGTTGTCGGTGTCTAG

*Barcode (BC) for round 3:*
CGATTCCGTTTGTAGTCGTC

*Barcode (BC) for round 4:*
AGTGCGTCATTGCAAGTAGT

*Barcode (BC) for round 5:*
GTCTTGTGTCCCAGTTACCA

#### B. Droplet microfluidics

Chip and emulsions chamber manufacture

Masters for the microfluidic chips were fabricated via soft lithography by Elliot Medcalf and Dr. Tomasz Kaminski. Single-use chips were cast by mixing PDMS with curing agent at a 10:1 ratio (Silicone Elastomer Kit Sylgard 184) and pouring the mixture onto the master. The liquid PDMS was degassed under vacuum and cured overnight at 65 °C. Hardened chips were cut to size, and 1 mm diameter holes were punched using a biopsy punch (Medline Industries). After plasma treatment of both PDMS and clean glass slides (Diener Femto), chips were bonded to the slides. All channels, except for electrode channels, were treated with 1% (v/v) trichloro(1H,1H,2H,2H-perfluorooctyl)silane dissolved in fluorinated oil (HFE-7500, Novec). Treated chips were incubated overnight at 65 °C. Emulsion chambers for droplet incubation and storage were prepared from standard 0.5 mL reaction tubes. Holes (1 mm diameter) were punched into one side and the bottom of each tube, and 0.5 × 1 mm PTFE tubing (Bola) was attached. The modified tubes were glued upside down onto glass slides.

Composition of the microfluidic rig

To prevent clogging during droplet microfluidics, all solutions were filtered prior to use with Merck Millex-GV filters, Whatman GD/X 13 filters, or Sartorius Minisart syringe filters. The oil phase consisted of HFE-7500 supplemented with 2% (w/v) 008-fluorosurfactant (RAN). The microfluidics setup comprised a syringe pump system (Nemesys) connected via tubing to PDMS chips, which were positioned under a conventional confocal microscope coupled to a high-speed camera (Phantom, Vision Research). Electronics for pico-injection and droplet sorting included a function generator (TG200, 20 MHz, TTi), a pulse generator (TGP110M, 10 MHz, TTi), and an amplifier (TRek, Model 601C).

Library transformation and expression

Before screening, the library was transformed into the expression host Bl21-DE3 (NEB). Transformation efficiency was confirmed by colony counting to reach sufficient oversampling of the library. Glycerol stocks with $OD_{600}$ of 1 were prepared and stored at -80°C. Library expression was conducted in bulk by inoculating one glycerol stock (V = 1 mL) in 5 mL LB media plus 50 µg/ml kanamycin. The culture was grown to $OD_{600}$ of 0.5 at 37 °C, 180 rpm. After induction with 0.2 mM IPTG, expression was performed for 18 h at 20 °C, 180 rpm.

Droplet generation, pico-injection and sorting for Absorbance Activated Droplet Sorting (AADS)

Droplets were generated at 1000 Hz using a flow-focussing device. Herein, cells were encapsulated together with substrate and lysis reagents in oil. To reach confident single-cell encapsulation according to Poisson-distribution, a $\lambda$ of 0.25 was used. After encapsulation, the cell suspension had an $OD_{600}$ of 0.00294 in 100 mM Tris/HCl pH 8 buffer with 20% Percoll (Sigma-Aldrich). Final concentrations of substrates and lysis reagents were 10 mM carbonyl substrate and 20 mM amine substrate, or 10 mM imine only, as well as 5 mM NADPH, 0.1 mg/mL polymyxin B, 1 µL/mL r-lysozyme, and 0.5 mM EDTA in 100 mM Tris/HCl pH 8 buffer. Flowrates were adjusted to yield droplets of 170 pl volume. Droplets were incubated off-chip and incubation times differed depending on the substrate. Pico-injection was performed at 350 Hz using a pico-injection device. Pico-injection solution contained at a final concentration 5 mM WST-1 (NBS biologicals), 1.67 µg/mL 1-methoxy-5-methylphenazinium methyl sulfate (mPMS, TCI), and 2 mM tartrazine in in Tris/HCl pH 8 buffer. Flowrates were adjusted to inject 85 pl volume per droplet. Voltages between 1.4 – 1.8 were used to allow injection and exclude droplet fusion. Absorbance measurements and sorting were performed at 150 Hz using a AADS device as well as custom-made Arduino and LabVIEW scripts. Voltages between 4 – 6 were used and backpressures were carefully adjusted using the outlet tubing of the positive and the negative channel.

Plasmid recovery after screening

Positive droplets were collected in a DNA LoBind tube (Eppendorf). The used tubing was carefully flushed with 2 ng/µl Salmon sperm DNA in DNAse free water after sorting. The emulsion was broken using 1H,1H,2H-2H-Perfluoro-octanol which was added to the collection tube in a 1:2 (v/v) ratio compared to the oil accumulated during sorting (or by using 100 µL 1H,1H,2H-2H-Perfluoro-octanol if no oil was collected in the tube of the positive channel). DNA was extracted three times by adding 2 ng/µl Salmon sperm DNA in DNAse free water, vortexing, and centrifugation. The aqueous phases were pooled and concentrated using a Monarch PCR and DNA clean-up kit from NEB. Final DNA was eluted in 6 µl nuclease-free water and transformed into 50 µL electrocompetent *E. coli* (E.cloni 10G Elite, Cambridge Bioscience).

#### C. Generation of the dETM dataset

Oxford Nanopore sequencing

Nanopore sequencing was performed as described previously(*4*). After each round of directed evolution, sorted variants were recovered by transformation into *E. coli*. Plasmid DNA for sequencing was prepared as follows: 10 mL of LB medium was added to the agar plate containing grown colonies, which were then scraped off with a sterile L-scraper. The resulting cell suspension was collected, cells were pelleted by centrifugation at 4000g, and plasmid DNA was isolated using the GeneJET miniprep kit according to the manufacturer's instructions. The plasmid was linearized by restriction digestion, and the resulting fragment was purified via agarose gel electrophoresis followed by SPRIselect bead cleanup. A total of 300–500 fmol of DNA was sequenced on a MinION R10.4 flow cell (Oxford Nanopore Technologies) using the ligation sequencing kit (V18), following the manufacturer's protocol. During bioinformatic downstream processing using the UMIC-seq pipeline(*3*), demultiplexing of the five rounds of directed evolution was performed using the round-specific barcode (BC).

Droplet microfluidics-based enrichment assay

Recovered DNA from each round of directed evolution was pooled and transformed into the expression host Bl21-DE3 (NEB). Droplet experiments were performed using AADS(*5*) with a 30 min off-chip incubation time. For sorting, a stringent gate (top 0.15% of droplets) and a lenient gate (top 1.5% of droplets) were chosen. For each gate droplet experiments were performed in triplicate, restarting from different stocks of the input libraries. Input and output libraries were kept and subjected to Illumina NGS.

NGS Illumina sequencing

Illumina sequencing was performed as described previously(*4*). Input and output libraries of the droplet-based enrichment assay were sequenced. Primers containing Illumina adapters and indices in their overhangs were used to amplify the round-specific barcode (BC) and UMI region from the recovered DNA. Output and input libraries from all microfluidics runs were multiplexed using primer pairs with distinct NGS index pairs. PCR mixes were prepared using the NEBNext Q5 mix (NEB) according to the manufacturer's instructions. PCR cycle numbers were optimized by amplifying 15% of the recovered DNA and performing qPCR, with the cycle number determined at 10% of the maximum fluorescence. Subsequent amplifications used 12–24 cycles. DNA was purified with SPRIselect beads (Beckman Coulter) at a 0.7:1 beads-to-sample volume ratio to remove primer dimers. DNA concentration was quantified using the NEBNext library quantification kit (NEB) and assessed by capillary electrophoresis (Bioanalyzer, Agilent), following the respective manufacturers' protocols. Depending on DNA quality and residual primer dimers, SPRIselect purification and quality control were repeated until acceptable purity was obtained. All purified input and output libraries were pooled in nuclease-free water to a final concentration of 0.2 nmol and analysed by the sequencing facility of the Department of Biochemistry (University of Cambridge). Illumina MiSeq libraries were prepared according to the manufacturer's instructions. Sequencing was performed on an Illumina MiSeq platform using a 2 × 150 bp run with 20% PhiX spike-in (500 cycles, 7.5 Gb).

Generation of fitness scores

Illumina sequencing was performed as described previously(*4*). A lost input library for the second screening replicate of the stringent sort was substituted with the second screening replicate of the lenient sort. A variant identifier file (UMI-identity dictionary) for downstream data generation was generated by custom python scripts. Fitness scores were calculated from the variant identifier file and the raw NGS paired end reads using the DiMSum pipeline[45] with the following parameters for the lenient and stringent sorts:

--gzipped = T --stranded = T --paired = T --barcodeErrorRate = 0.1 --cutadapt5First = ATTCGATGGTGTCCGGGATCTCGACGCTCTCCCTTAGATGC --cutadapt5Second = AGTCGTATTAATTTCCTAATGCAGGAGTCGCATATAGTGAGTCG --cutadaptMinLength = 20 --cutadaptErrorRate = 0.6 --cutadaptOverlap = 3 --vsearchMinQual = 10 –vsearchMaxQual = 41 --vsearchMaxee = 0.5 --vsearchMinovlen = 10 --wildtypeSequence = *Sr*IRED$_{wt}$ (Supplementary Table 1) --reverseComplement = F --sequenceType = auto --mutagenesisType = random --transLibrary = F --transLibraryReverseComplement = F --bayesianDoubleFitness = F --bayesianDoubleFitnessLamD = 0.025 --fitnessMinInputCountAll = 0 --fitnessMinInputCountAny = 0 --fitnessMinOutputCountAll = 0 --fitnessMinOutputCountAny = 0 --fitnessHighConfidenceCount = 10 --fitnessDoubleHighConfidenceCount = 50 --fitnessNormalise = T --fitnessErrorModel = T --indels = all --maxSubstitutions = 100 --mixedSubstitutions = T --retainIntermediateFiles = T --retainedReplicates = all --startStage = 1 --stopStage = 5.

Fitness scores ($F_i$) as defined by the DiMSum pipeline are the natural logarithm of enrichment score $W_i$ defined by the enrichment (measured with NGS counts $N$) of a variant in the output library ($i$) with respect to the input library, standardized to the enrichment of wildtype ($wt$):

$$F_i = \ln(W_i) = \ln(\frac{\frac{N_i^{output}}{N_i^{input}}}{\frac{N_{wt}^{output}}{N_{wt}^{input}}})$$

Output fitness scores obtained for the stringent and lenient sorts were subjected to several quality control thresholds to obtain high-confidence and low-confidence fitness scores (details are provided in **Figure S4**).

Generation of $k_{cat}^{app}$ from fitness scores

$k_{cat}^{app}$ was estimated from the DMS-derived fitness values by fitting an exponential relationship between experimentally measured turnover numbers and the corresponding variant fitness scores. The model fit

$$k_{cat} = a * e^{b*fitness}$$

Only variants with both measured $k_{cat}$ and fitness_final values, and with kcat > 0, were included in the fitting process. The fitted parameters were then applied to all variants in the fitness dataset to calculate $k_{cat}^{app}$ from fitness_final. Uncertainty in $k_{cat}^{app}$ was propagated from the reported fitness error (sigma_final) using the derivative of the exponential mapping, such that

$$k_{cat\_app\_error} = |b| * k_{cat(app)} * sigma$$

This produced a transformed dataset in which each variant retained its original fitness values and associated errors, together with the inferred $k_{cat}^{app}$ and propagated $k_{cat}^{app}$ error.

#### D. Protein work

<u>*Sr*IRED expression</u>

The plasmid encoding for *Sr*IRED was transformed into the expression host Bl21-DE3 (NEB). A pre-culture in 5 mL LB media plus 50 µg/ml kanamycin was grown over night (37 °C, 200 rpm). An expression culture was inoculated and grown to $OD_{600}$ of 0.8 at 37 °C, 180 rpm. After induction with 1 mM IPTG, expression was performed over night at 20 °C, 180 rpm. Cells were harvested by centrifugation (4000 g) and resuspended in lysis buffer (100 mM phosphate pH 7, 100 mM NaCl, 1 mM DTT, Roche complete protease inhibitor tablet).

<u>*Sr*IRED purification</u>

Cell lysis was performed using sonication for 10 min (Vibracell, 40% amplitude, 1 s on – 1 s off). The cell lysate was clarified at 14,000 g for 45 min. For purification, lysate supernatant was incubated with NiNTA agarose (NeoBiotech) on a roll mixer at 4 °C for 1 h. Subsequently, the NiNTA agarose was transferred into a gravity column with frit. Three wash steps were performed with 15 mL wash buffer (100 mM phosphate pH7, 100 mM NaCl, 1 mM DTT, 30 mM imidazole). The protein was eluted using 10 mL elution buffer (100 mM phosphate pH7, 100 mM NaCl, 1mM DTT, 300 mM imidazole) and rebuffered into storage buffer (100 mM phosphate pH7, 100 mM NaCl, 1 mM DTT, 10% glycerol) using amicon ultra centrifugal filters (10,000 NMWL, Merck). Aliquots were frozen with liquid nitrogen and stored at -80°C.

#### E. Enzyme activity assays

<u>Secondary assays after microfluidics screenings in lysate</u>

After microfluidics sorting, the recovered library was retransformed into chemically competent Bl21(DE3) cells (NEB). 90 random colonies were picked into 1 mL LB media plus 50 µg/ml kanamycin in deep-well plates (Greiner). These were grown over night at 37 °C, 750 rpm. Expression cultures were set up by diluting 30 µl of this preculture in 900 µl LB medium plus kanamycin in duplicates. After induction at $OD_{600}$ of 0.4 – 0.6 with 0.2 mM IPTG, the protein was expressed for 18 h at 20 °C, 750 rpm. Cells were harvested by centrifugation (45 min, 4000g) and lysed for 1 h (20 °C, 750 rpm) using 100 – 200 µl lysis buffer per well (100 mM Tris pH 8, 4 mg/ml egg white lysozyme, 0.5 mM EDTA, 0.75 mg/ml polymyxin B). Lysate was clarified by centrifugation (45 min, 4000g, 4°C) and supernatant was used for subsequent activity assays. IRED activity was determined as initial rates of NADPH depletion at 340 nm in a plate reader (Spectramax 190, Molecular Devices). Assays were performed in Nunc 96-well plates (Thermo) with 10 mM carbonyl substrate and 20 mM amine, or 10 mM imine only, as well as 0.5 mM NADPH in 100mM Tris-HCl buffer at pH 8. Activities were normalised on the activity of the wild-type enzyme within the same experiment.

Michaelis-Menten kinetics of *Sr*IRED

Kinetic constants of *Sr*IRED variants were determined in 100 mM Tris pH 8.0 with saturating concentrations of two co-substrates and varying concentration of the third co-substrate. The reactions were run in transparent Nunc 96-well plates (ThermoFisher) and absorbance of NADPH was monitored to derive enzyme rates (Spectramax 190, Molecular Devices). Between 0.0015 and 0.05 g/L enzyme was added to initiate the reaction, and measurements were carried out at 20 °C. Initial rates were measured in three replicate wells and Michaelis-Menten curves were fitted using Prism.

#### F. Bioinformatics

Network graph generation

A distance matrix was generated for the unique variants derived from Nanopore sequencing. This distance matrix was converted into a network graph in gml format using the Python package NetworkX(*6*). Visualisation was performed using Cytoscape(*7*) with the “Prefuse Force Directed Layout”.

Fitness landscape generation

The traversed fitness landscape of the directed evolution campaign was generated based on a network graph obtained from the fitness-labelled dETM dataset. This network graph provided the 2D x-y-coordinates of the fitness landscape. Fitness values were added as z-axis and a smooth surface was generated through linear interpolation, leading to a 3D fitness landscape.

Analysis of epistasis using fitness scores

To calculate epistatic interactions for more higher-order variants, the available dETM data was complemented with external data of single mutation fitness derived from a deep mutational scan conducted on the same *Sr*IRED wildtype epPCR library by Gantz *et al.* (2026)[9]. Epistatic scores ($\varepsilon$) were quantified from single mutants as base scenario by deriving the difference between higher-order fitness ($F_{abc}$) single mutant fitness ($F_a + F_b + F_c$):

$$\varepsilon_{from\ singles} = \ln(W_{abc}) - \ln(W_a * W_b * W_c) = F_{abc} - (F_a + F_b + F_c)$$

Additionally, epistatic scores were quantified based on the respective parental higher-order mutant ($F_{ab}$) and the added single mutant ($F_c$):

$$\varepsilon_{from\ higher-order\ mutants} = \ln(W_{abc}) - \ln(W_{ab} * W_c) = F_{abc} - (F_{ab} + F_c)$$

Co-evolution analysis

For co-evolution analysis, the dETM dataset was filtered for each the founder mutations of interest. For these smaller datasets, amino acid mutation type and frequency per position was calculated.

Population dynamics analysis

Paired-end NGS reads of the five input library replicates were used to quantify the fraction of each variant in the population throughout the five rounds of evolution. The NGS reads were demultiplexed based on the round-specific barcode (BC) which included the merging of paired-end reads using NGmerge(*8*) with default settings and demultiplexing using the UMIC-seq helper script(*3*) with the following settings:

demultiplex --threshs 22 22, using the following fasta sequences: >round1 CCGGCGTAGAGGATCGAGA >round2 AAGTAGTTGTCGGTGTCTAG >round3 CGATTCCGTTTGTAGTCGTC >round4 AGTGCGTCATTGCAAGTAG >round5 GTCTTGTGTCCCAGTTACCA.

After demultiplexing and to obtain the raw NGS counts per round, these files were parsed through the DiMSum pipeline(*9*) to recover the total number of NGS counts per variant using the UMI-identity dictionary and applying the same settings as described above with a few modifications:

--cutadapt5First = ATTCGATGGTGTCCGGGATCTCGACGCTCTCCCTTAGATGC --cutadapt5Second = GAGTCGTATTAATTTCCTAATGCAGGAGTCGCATATAGTGAGTCG --paired = F.

Files outputted by DiMSum were only used to extract the number of counts per variant (in quintuplicate measurements) for each respective round of evolution. NGS counts for round 0 (complete naive starting library) were obtained from the source data from Gantz et al. (2024)(*4*), which used the same *Sr*IRED wildtype epPCR library.

Evolutionary history analysis

The closest 5,000 homologues of *Sr*IRED were retrieved through a BLAST search. This dataset was used for evolutionary history analysis. A phylogenetic tree was generated from an MSA which was obtained via MAFFT(*10*). The tree was built using FASTTREE(*11*) and visualized as a cladogram using iTOL(*12*).

Mutation hotspot analysis

The entire dataset of unique variants derived from Nanopore sequencing was used to analyse each position of the *Sr*IRED sequence for mutations. A stacked bar plot was generated reporting on amino acid mutation type and frequency per position.

Campaign fitness prediction

We benchmarked in silico models on the fitness landscape by asking whether each method predicted the direction of experimentally measured activity changes along local mutational edges. Variants were taken from the low and high-confidence fitness dataset. The low-confidence outlier A271T was excluded. Experimental fitness values were converted to apparent catalytic activity as $k_{cat}^{(app)} = 6.2\ e^{0.64*fitness\_final}$.

A genotype graph was constructed with variants as nodes. Edges connected pairs of variants differing by one amino-acid substitution, with the lower-order genotype treated as the source and the higher-order genotype as the target. Analyses were restricted to the WT-connected component, yielding 984 directed one-step mutational edges. The experimental sign for each edge was defined as the sign of the change in $k_{cat}^{app}$ from source to target.

Each method assigned one score to each variant. ESM2 (esm2_t33_650M_UR50D(*13*)) and ESM-C (esmc_300m(*14*)) were evaluated on unmasked variant sequences; the mean log probability of the observed residues was used as the sequence-plausibility score(*15*). MSA conservation scores were computed from the aligned homologous sequence set by summing, across mutated sites, the log-frequency change from WT residue to mutant residue. EVmutation was fit from the same MSA using plmc, and variants were scored by their epistatic Hamiltonian change(*16*). GEMME was run with the official web implementation on the query-column-trimmed MSA, and the evolCombi score was used(*17*).

ThermoMPNN(*18*) was run in site-saturation mode on the SrIRED crystal structure 5OCM(*19*). Chain B of 5OCM was parsed with the ThermoMPNN/ProteinMPNN structure featurization utilities, and all 20 amino-acid identities were evaluated at each resolved residue position. This produced a single-mutant prediction table containing 5,800 entries, with columns specifying position, WT residue, substituted residue, chain, and predicted ΔΔG. To score assayed multi-mutant variants we summed the predicted ΔΔG values across substitutions in the variant.

We also evaluated supervised sequence-to-activity models. A MULTI-evolve-style fully connected neural network (FCN) was trained with the used one-hot featurizer(*20*) to predict log-transformed $k_{cat}^{app}$. EVOLVEpro's random forest regressors were trained either on one-hot sequence encodings or on ESM2-650M mean-pooled sequence embeddings(*21*). These supervised models were trained only on WT, single-mutant and double-mutant variants and then used to score all other variants. For edge-level evaluation, supervised models were assessed only on edges that did not touch variants with mutational load under two amino acid mutations, thereby excluding training-neighbourhood edges.

For each model and each scored edge, the predicted direction was defined as the sign of the model-score difference between target and source. Pairwise agreement between two methods was calculated as the fraction of shared scored edges for which the two predicted signs were identical. Agreement with experiment was calculated analogously using the measured sign of $\Delta\ k_{cat}^{(app)}$.

ESM2 'holey landscape' assessment

Variant naturalness was estimated using ESM2 (esm2_t33_650M_UR50D) in single-pass mode(*15*). Full-length variant sequences from the high-confidence dETM dataset were passed through the model without masking, and the log probability assigned to the observed amino acid at each residue was extracted from the output logits. Residue log probabilities were averaged across the sequence and converted to a pseudo-perplexity score, $PPL^* = e^{-\text{mean log probability}}$, where lower values indicate higher model plausibility.

Each variant was assigned to the first campaign round in which it was observed. To generate a round-matched null, we defined, for each round, the pool of unique substitutions observed among variants present in that round. For every observed variant, five random null genotypes were sampled from the corresponding round-specific substitution pool with the same mutational load as the observed genotype. Samples containing repeated residue positions or substitutions inconsistent with the WT sequence were rejected and resampled.

Null sequences were scored with the same ESM2 procedure. For each round, we compared the median observed PPL* with the median PPL* of the corresponding round-matched null variants.

Sequence similarity networks

Dimensionality reduction of the dETM dataset was done using Levenshtein metric for distance calculation and UMAP (Uniform Manifold Approximation and Projection) or tSNE (t-distributed stochastic neighbour embedding) as visualisation tool. Code to reproduce the sequence similarity networks is available online at https://github.com/pauljannis/pySSN.

Simulating the greedy walk

To ask what the outcome of a traditional DE mutation would be fixing approach, we replayed the campaign and artificially fixed ‘winner mutants’ in each round of the campaign. After the first round, T241A was artificially fixed as the winner and alternative lineages from this founder followed. In each subsequent round, all descendants of a particular parent are evaluated. To constrain the decision tree to realistic experimental parameters, these descendants could not differ by more than two mutations per round and the trajectory they belong to must not end before the final round. The most fit descendant matching these criteria is fixed and the process repeats for the next round.

### Supplementary Figures

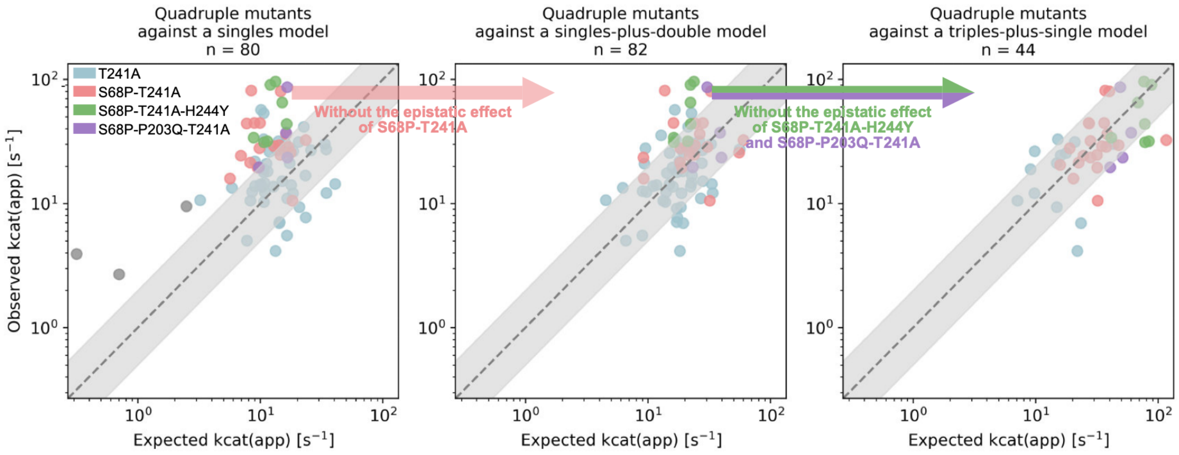

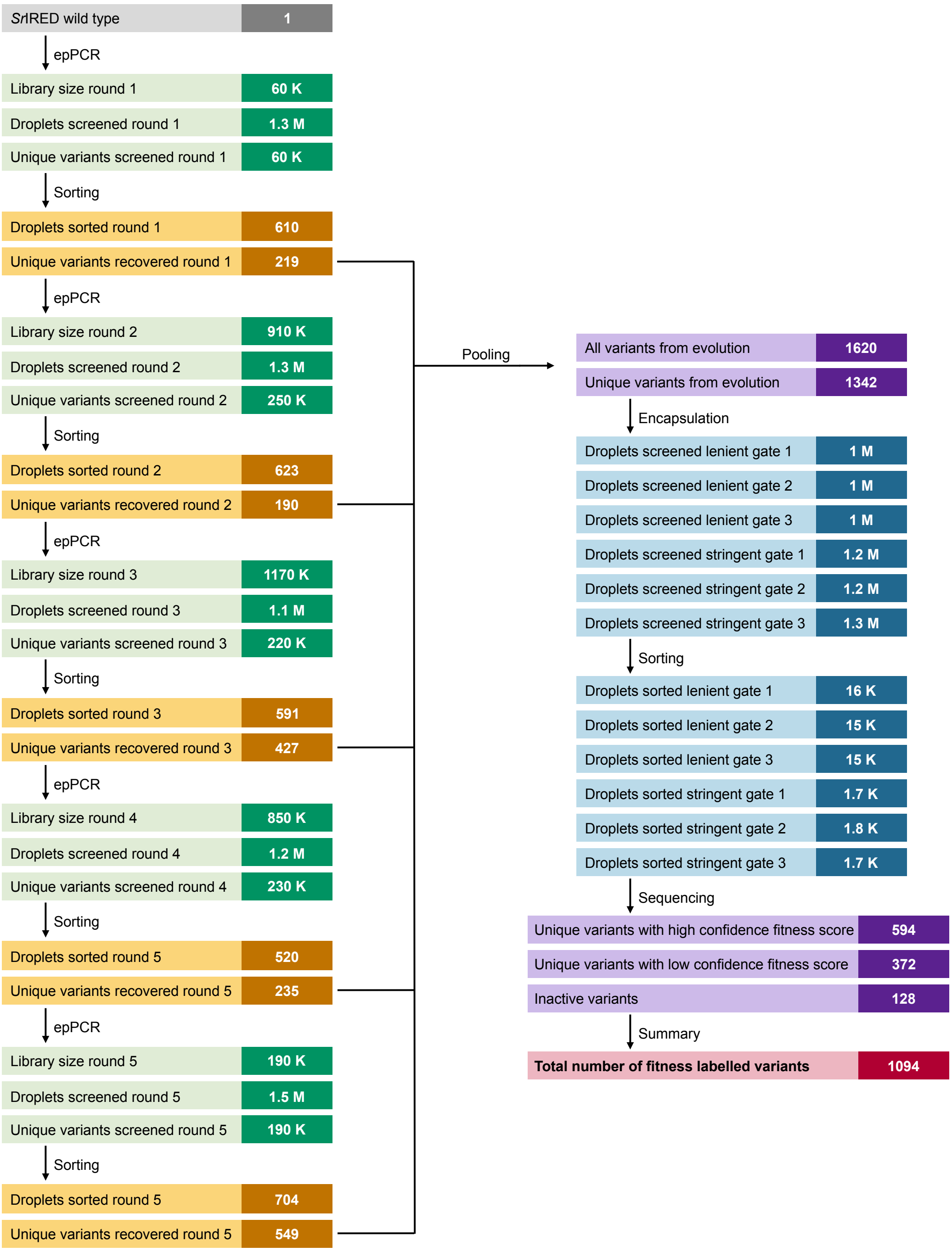

**Figure S1: Overview on numbers during evolution and fitness labelling.** The left-hand side summarises the directed evolution campaign and reports on library sizes, number of droplet-based experiments performed, and number of unique variants tested after considering droplet occupancy and oversampling. Discrepancies between screened droplets and recovered unique variants stem from variant replicates in the sorted droplets. The right-hand side summarises the fitness labelling which was performed with all recovered variants from the directed evolution campaign. Discrepancy between all variants from evolution and unique variants from evolution stem from variant replicates. In total, 950 K unique variants were tested during directed evolution in 6.4 M droplet-based experiments. Over five rounds, the best 1342 variants were selected. For fitness labelling, another 6.7 M droplet-based experiments were performed to resolve the fitness of 1094 variants.

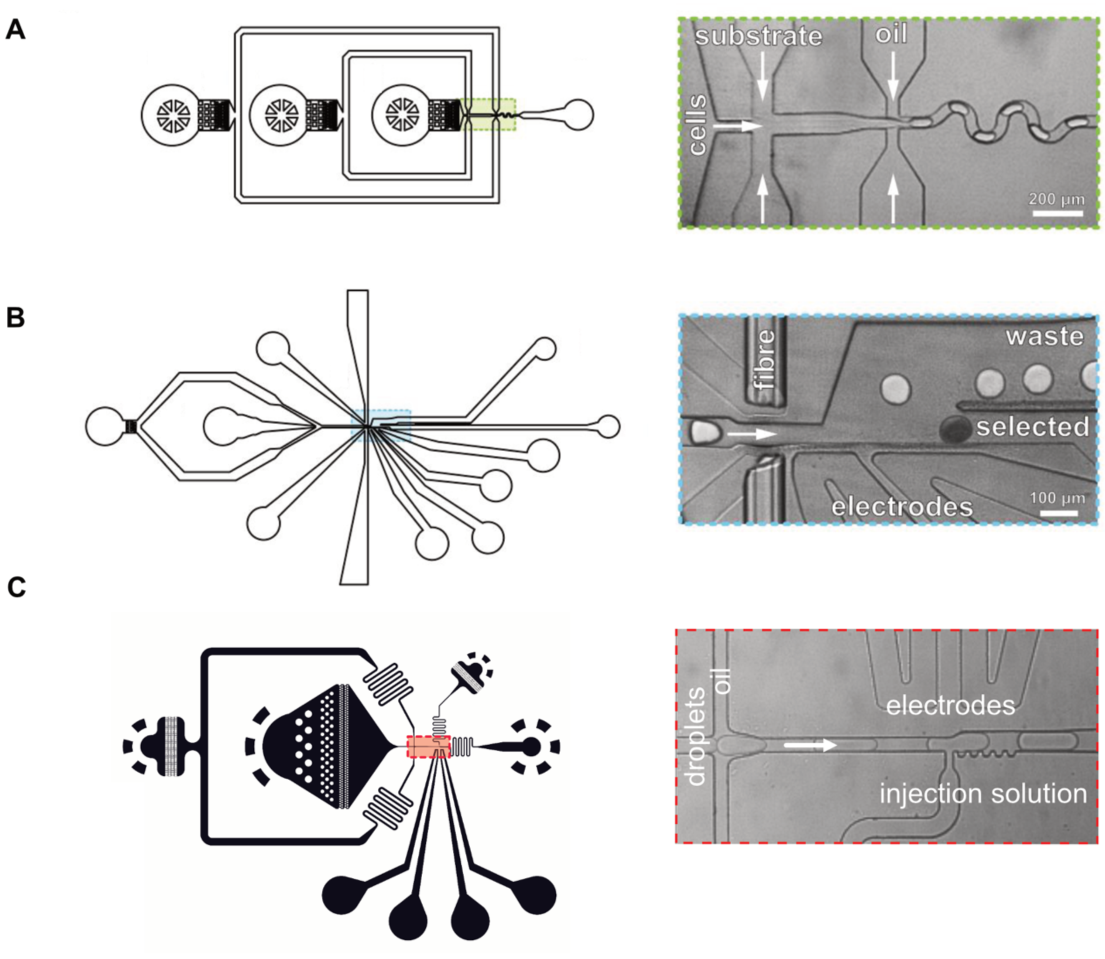

**Figure S2: Microfluidic chip designs.** All designs are deposited in CAD format on the DropBase repository (https://openwetware.org/wiki/DropBase:Devices). **(A)** Flow-focusing chip for droplet generation https://openwetware.org/wiki/DropBase:droplet_generation_3_inlets **(B)** Sorting chip https://openwetware.org/wiki/DropBase:AADS **(C)** Picoinjection chip https://openwetware.org/wiki/DropBase:Picoinjection_chip.

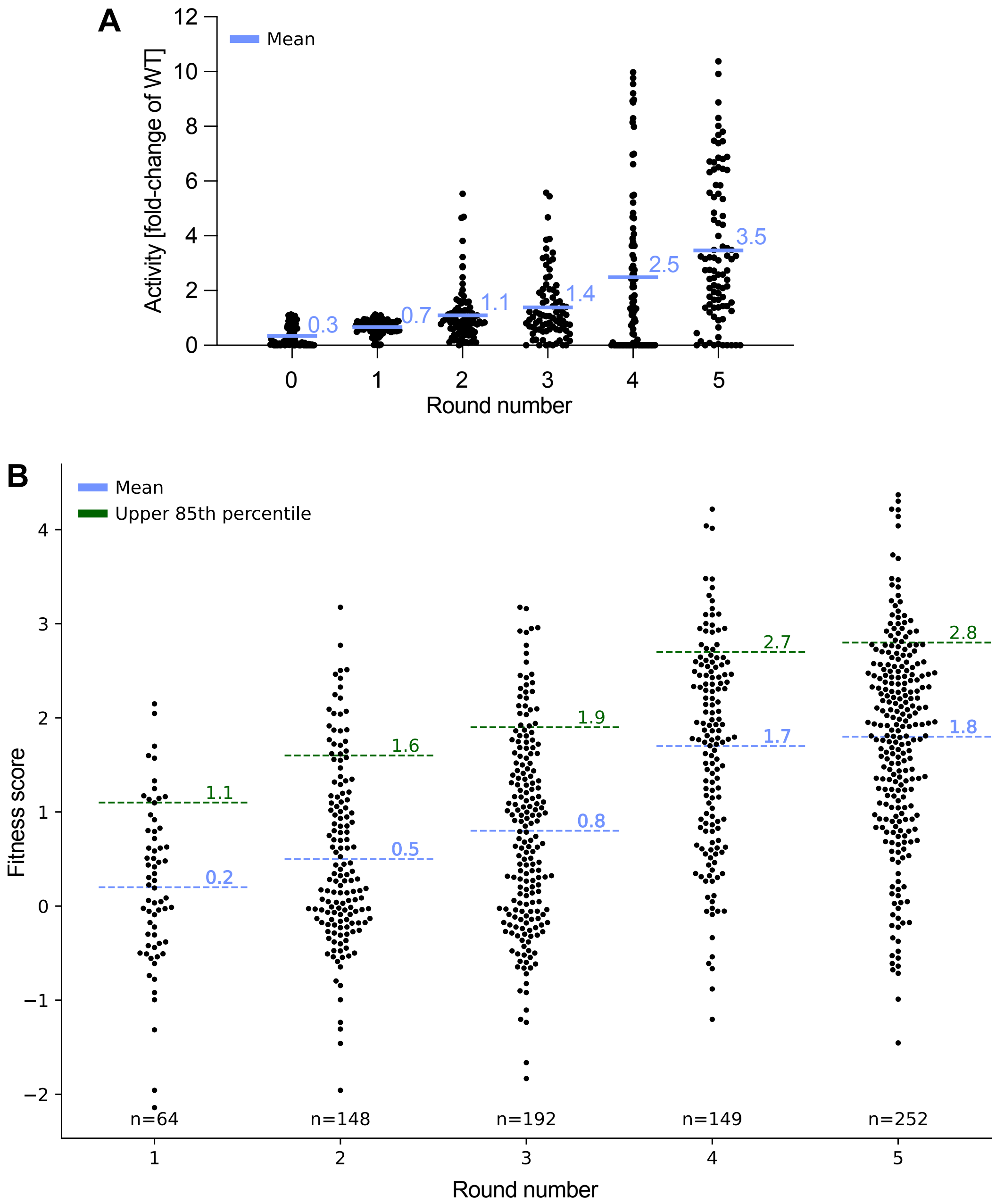

**Figure S3: Fitness improvements during directed evolution. (A)** Plate-based secondary assay for five rounds of directed evolution in lysate. 90 random variants from the first input library as well as from each sorted library after a round of directed evolution were grown in 96-well-plate format and expressed for 18 h at 20°C. Cells were lysed and activities of clarified diluted supernatants were determined as initial rates of NADPH depletion at 340 nm. Reactions were performed with 10 mM **1**, 20 mM **a**, and 0.5 mM NADPH in 100mM Tris-HCl buffer at pH 8. Activities were normalised on the activity of the wild type (WT). Average activity is indicated in blue. All results are shown as average of two technical replicates. **(B)** Bee swarm of all fitness labelled variants from the directed evolution campaign. After high-confidence fitness scores were generated, they were allocated to the rounds of evolution where the respective variants occurred. Mean of the population is shown in blue and upper 85th percentile in green.

**A**

| Gate | | Droplets screened [x $10^6$] | Droplets sorted | Incubation period [min] | AADS over-sampling (20% droplet occupancy) | NGS reads input [x $10^6$] | NGS depth input | NGS reads output [x $10^5$] | NGS depth output |
|---|---|---|---|---|---|---|---|---|---|
| | 1 | 1.03 | 15,560 | ~ 30 | 149 | 1.63 | 1,212 | 1.44 | 107 |
| Lenient | 2 | 1.03 | 15,435 | ~ 60 | 149 | 0.98 | 728 | 1.26 | 94 |
| | 3 | 1.02 | 15,230 | ~ 30 | 148 | 1.09 | 811 | 2.14 | 159 |
| | 1 | 1.23 | 1,665 | ~ 30 | 178 | 1.30 | 967 | 2.50 | 187 |
| Stringent | 2 | 1.24 | 1,761 | ~ 30 | 180 | 0 | 0 | 1.59 | 118 |
| | 3 | 1.27 | 1,676 | ~ 30 | 184 | 0.92 | 685 | 1.43 | 106 |

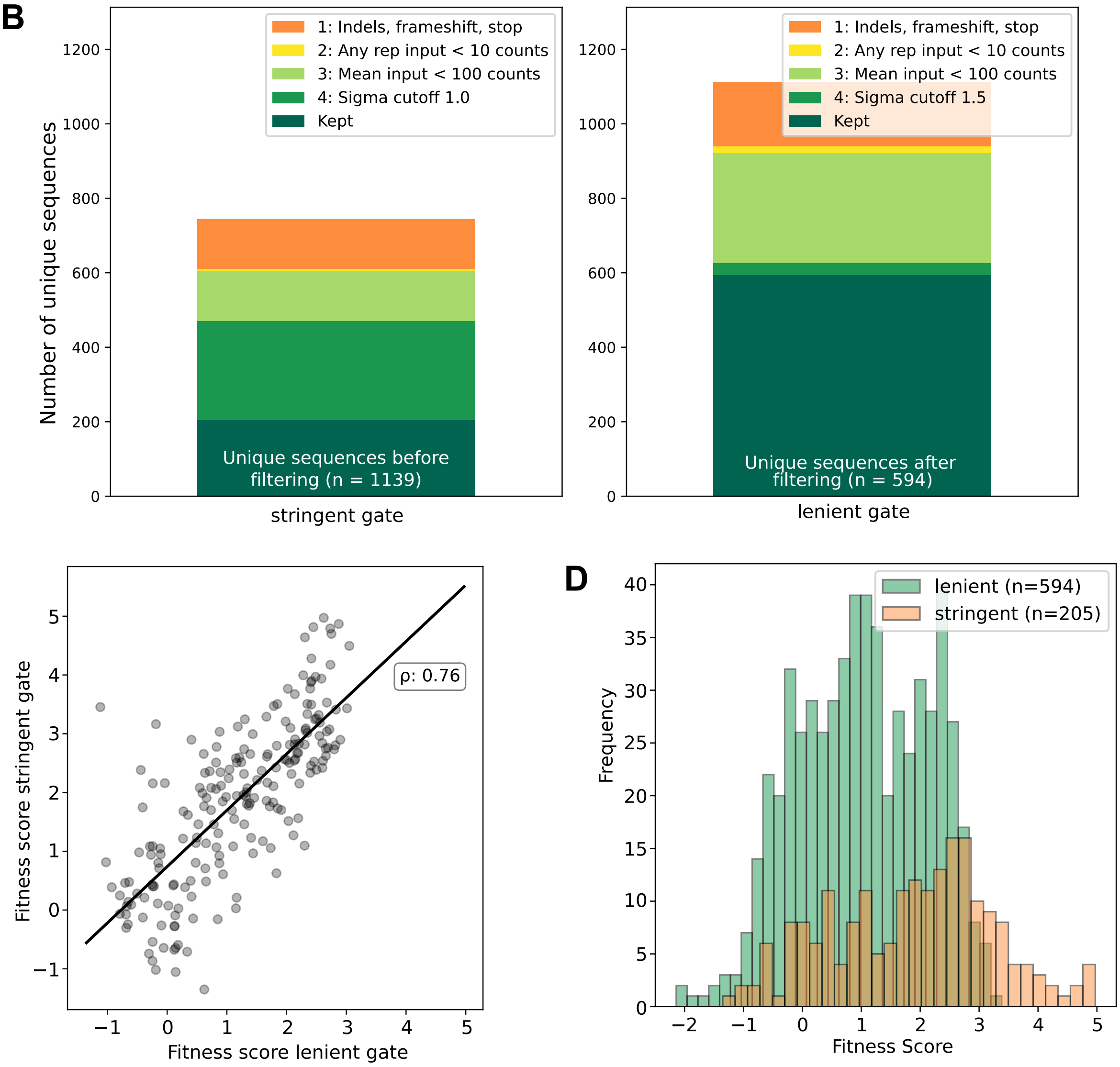

**Figure S4: Workflow for generating fitness scores. (A)** The number of screened and sorted droplets are depicted for each respective replicate in the lenient and stringent sorting gate. A short in-droplet incubation period was chosen to keep in the dynamic range of the assay. The AADS oversampling rate refers to how often each variant in the full naive library (1,342 unique variants) was encountered during AADS assuming that 20% of droplets were occupied by an *Sr*IRED variant using the Poisson distribution. Numbers of NGS reads in the input (before sorting, naive library) and the output (after sorting) libraries are depicted, and sequencing depth was calculated referring to the coverage of the input (3,040 unique UMI's) and output (estimated at ¼ of the input diversity) libraries by NGS sequencing. **(B)** Post-DiMSum filtering of lenient and stringent gates. Bar plots contain all unique sequences within the stringent and lenient datasets respectively, and it is indicated at which steps sequences were excluded during the post DiMSum filtering process. Filters included removing InDels, frameshifts and premature stop codons, removing sequences with too low input counts (as this is the main indicator for accurate fitness scores[13]), and removing sequences that constitute uncertainty outliers. Filter thresholds were different for lenient and stringent gates and are detailed within the figure panel. **(C)** Fitness score correlation between both gates shows that both gates report on the same variant property. The top right value denotes Pearson's r. **(D)** Histogram of fitness scores in the lenient and stringent gates shows different dynamic ranges for both gates with the lenient gate reporting better on low-fitness variants and the stringent gate reporting better on high-fitness variants.

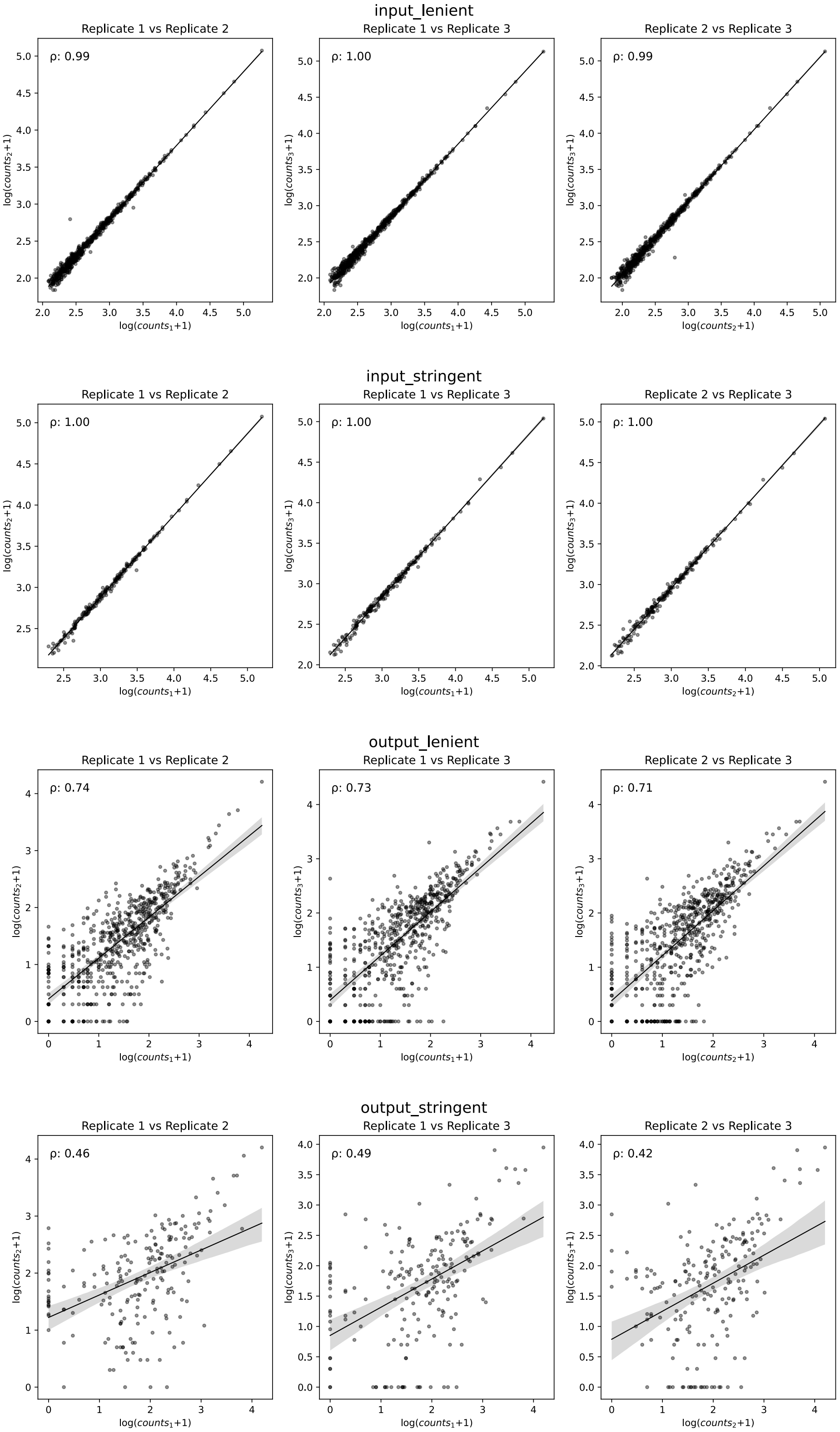

**Figure S5: Correlation between replicates of the enrichment assay used for fitness labelling.** Correlation plots between all replicates for all unique sequences which passed the post lrDMS filtering regime within each sorting threshold: lenient (n = 389) and stringent (n = 205).

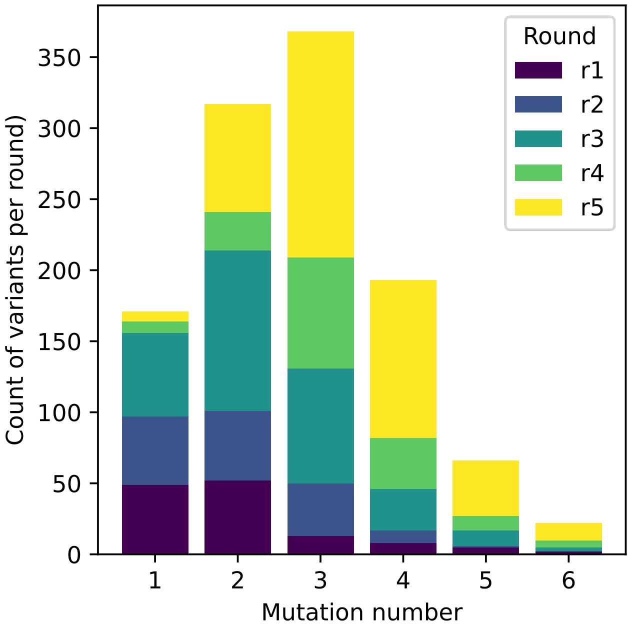

**Figure S6: Overview of the dETM dataset for the cyclohexanone and cyclopropylamine campaign.** Distribution of mutational load across all unique *Sr*IRED sequences which carry a high-confidence fitness score (594 variants).

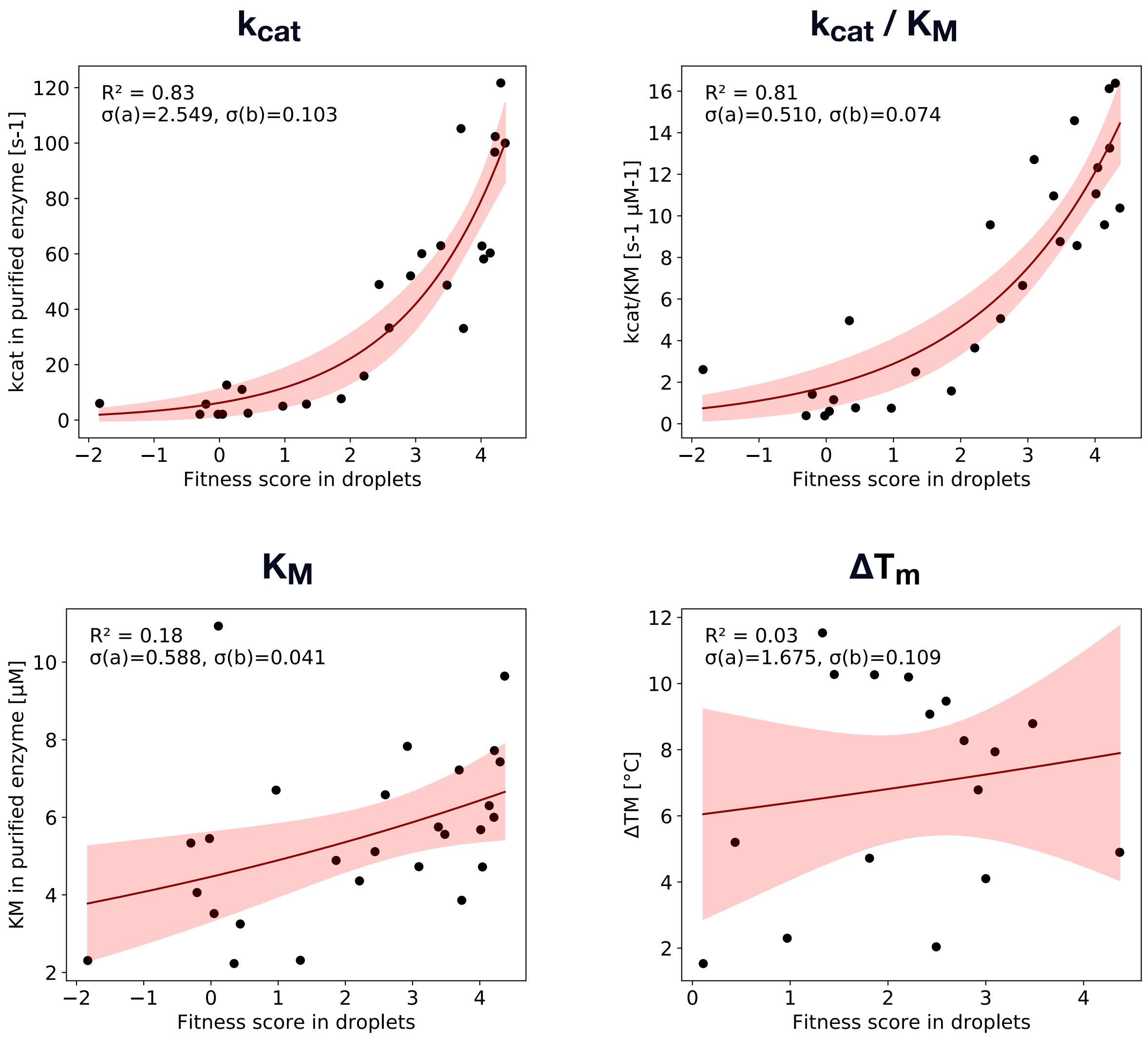

**Figure S7: Correlations between droplet-based fitness scores and biophysical as well as kinetic parameters.** Fitness scores derived from droplet experiments are cumulative labels reporting on total turnover of substrate in droplets. For 27 variants, fitness scores and biophysical as well as kinetic parameters ($T_M$, $K_M$, $k_{cat}/K_M$) are correlated using exponential fit.

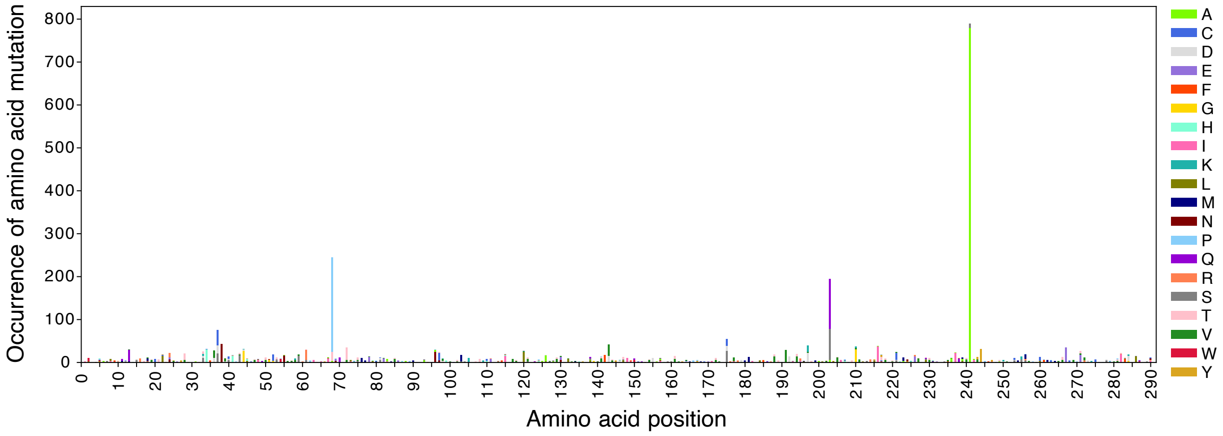

**Figure S8: Hotspot analysis after five rounds of directed evolution.** All 1342 unique variants obtained during directed evolution were analysed collectively to determine the mutation occurrence at each amino acid position of the *Sr*IRED sequence. In the entire dataset, position 241 was mutated in 58.9% of all sequences, position 68 was mutated in 18.3% of all sequences, and position 203 was mutated in 14.5% of all sequences.

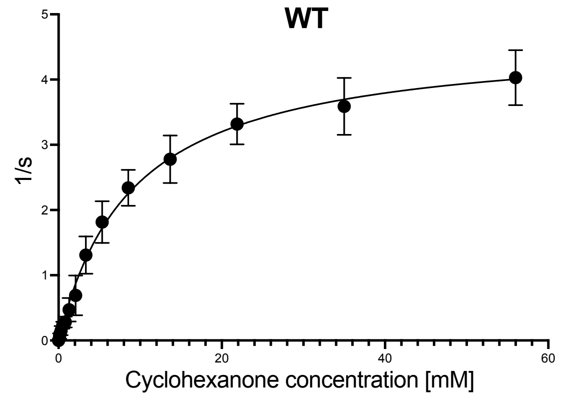
WT
1/s
Cyclohexanone concentration [mM]

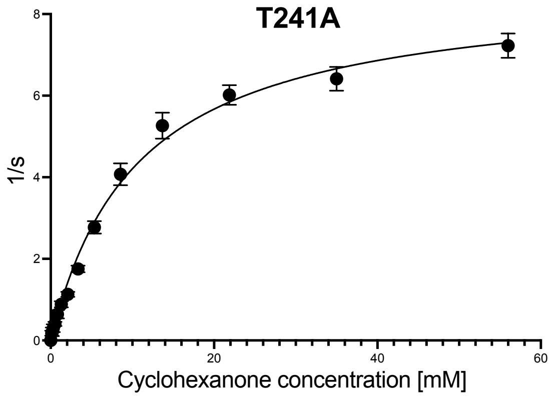
T241A
1/s
Cyclohexanone concentration [mM]

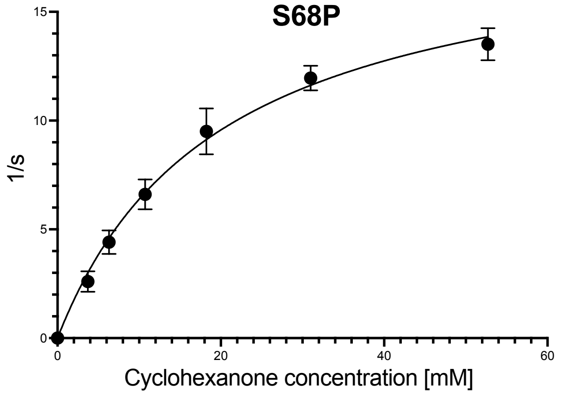
S68P
1/s
Cyclohexanone concentration [mM]

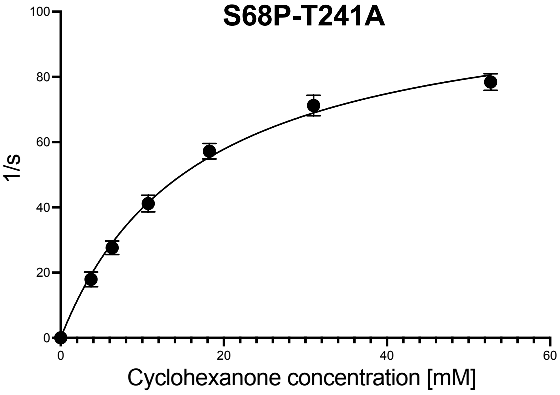
S68P-T241A
1/s
Cyclohexanone concentration [mM]

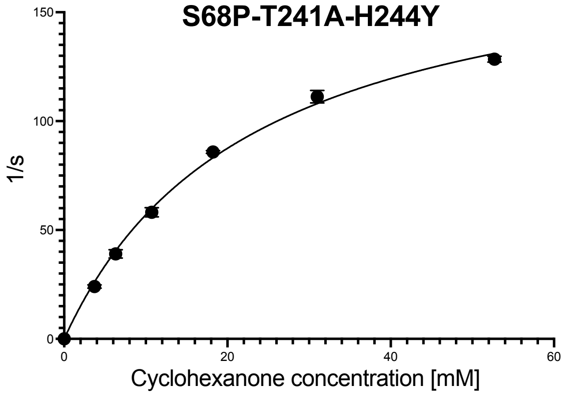
S68P-T241A-H244Y
1/s
Cyclohexanone concentration [mM]

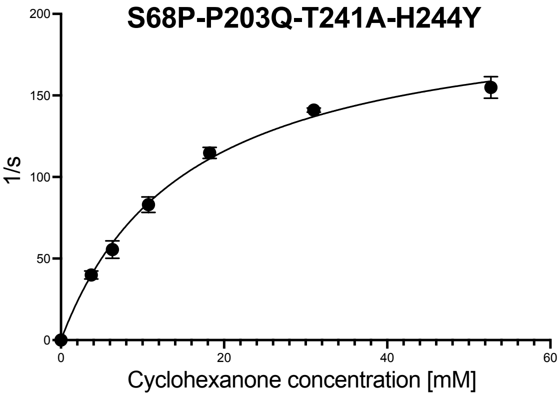
S68P-P203Q-T241A-H244Y
1/s
Cyclohexanone concentration [mM]

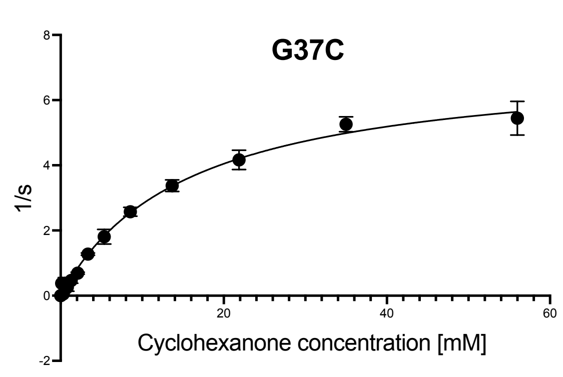
G37C
1/s
Cyclohexanone concentration [mM]

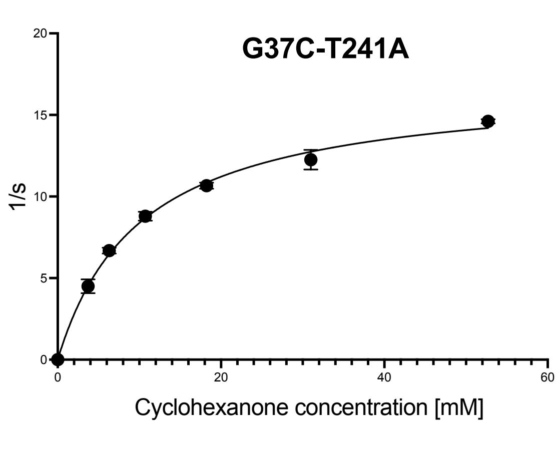
G37C-T241A
1/s
Cyclohexanone concentration [mM]

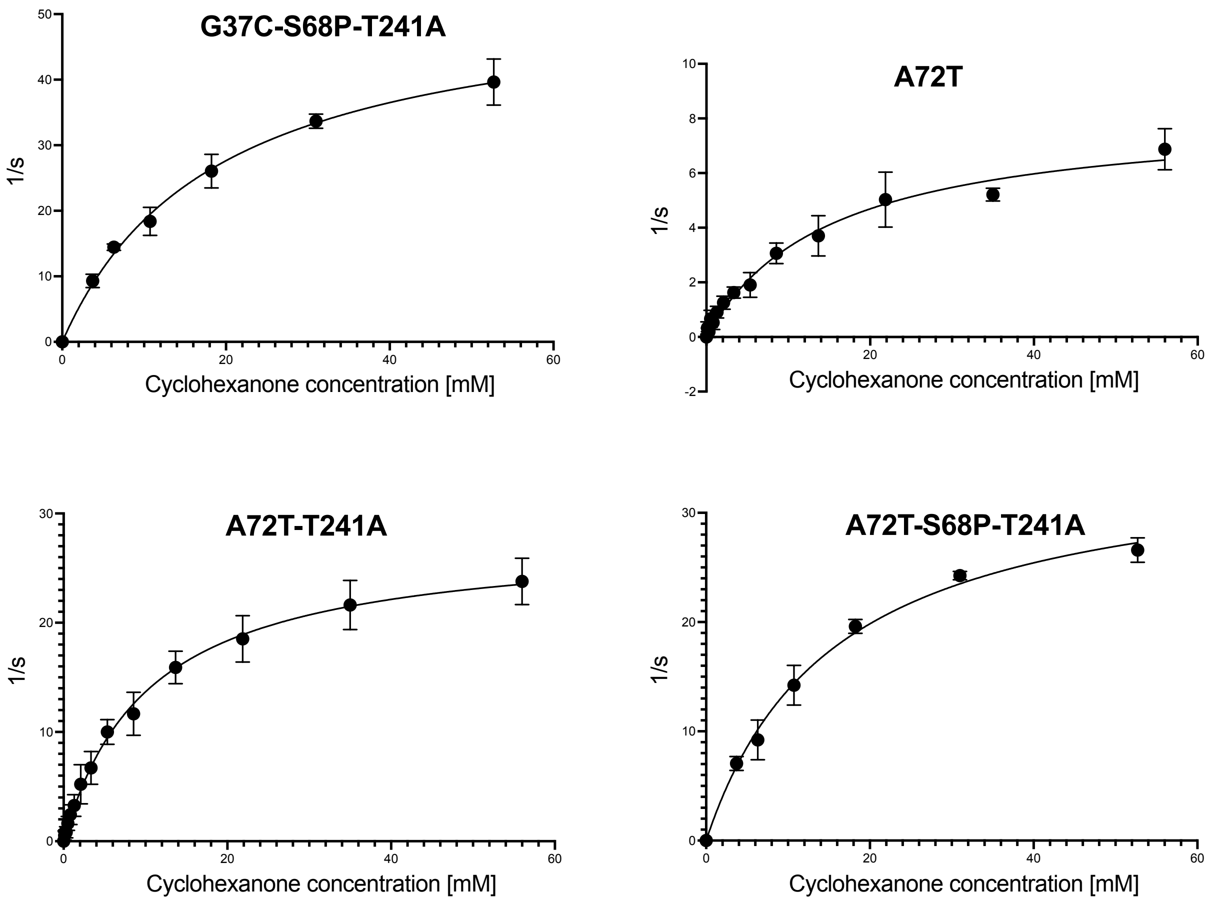

**Figure S9: Michaelis-Menten plots for cyclohexanone kinetics.** Kinetics were conducted with variable concentrations of cyclohexanone (0.1 - 52.7 mM) and a constant concentration of cyclopropylamine (75 mM) and NAPDH (1.5 mM) in Tris pH 8 with variable enzyme concentrations (from 0.0015 to 0.05 g/L) depending on the rate at T = 25°C. Initial rates were calculated based on NADPH depletion at 380 nm using a standard curve of NADPH at that wavelength. Summary of all Michaelis-Menten parameters is provided in **Error! Reference source not found.**.

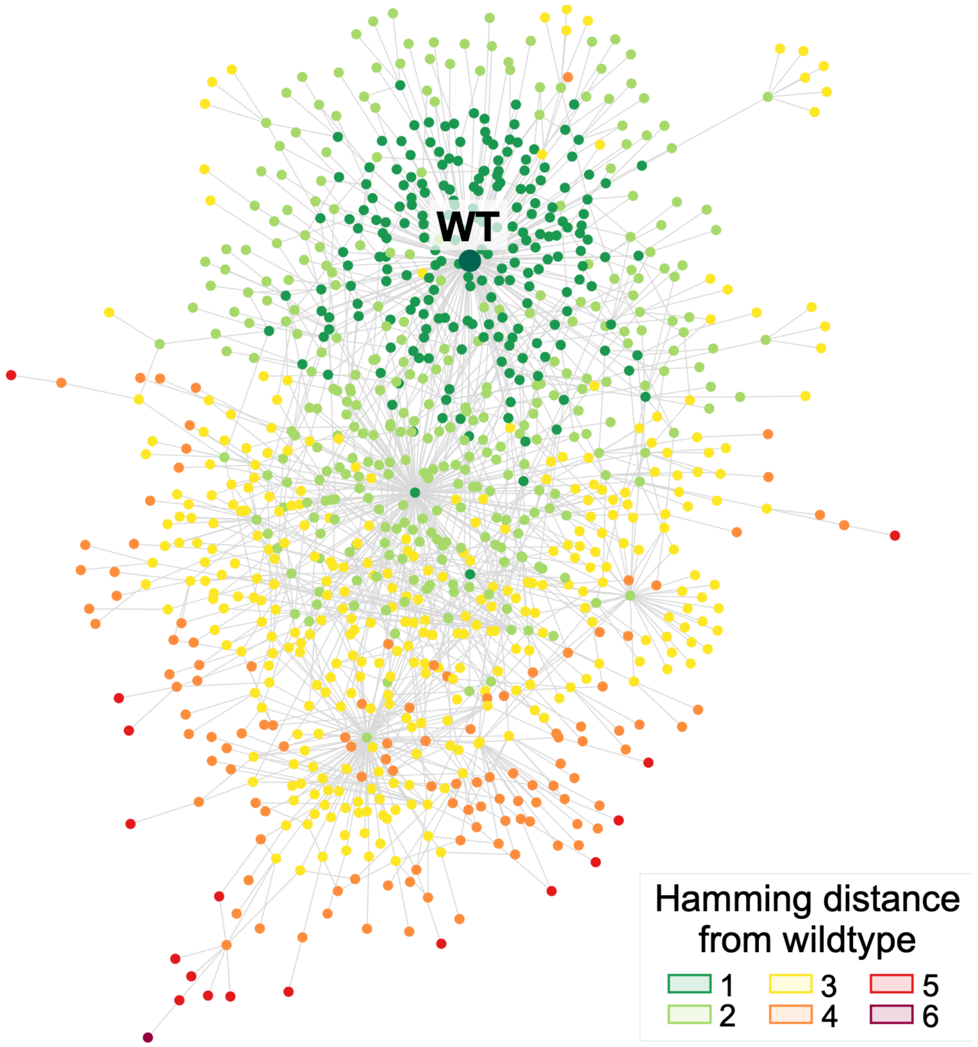

**Figure S10: Network graph coloured by mutation number.** Network graph of the recorded evolution campaign where each node represents one variant, and two nodes are connected by one edge if respective variants are one amino acid mutation away from each other (total of 990 nodes and 1346 edges). Wildtype (WT) is marked in dark green. Distances in the network graph correspond to Hamming distances from wildtype in the explored sequence space.

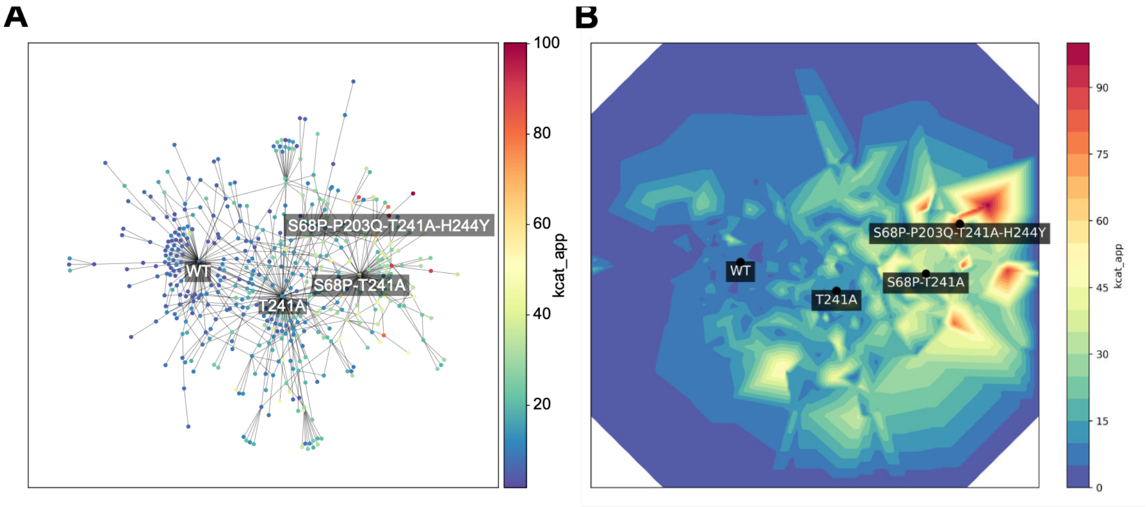

**Figure S11: Generation of the campaign's fitness landscape. (A)** Network graph generated with the 594 variants which carry a $k_{cat}^{app}$ label. **(B)** Contour gradient of the campaigns fitness landscape. $k_{cat}^{app}$ scores were used as z-axis on the previously generated network graph and displayed as contour gradient obtained through linear interpolation. Unmeasured space was assigned a $k_{cat}^{app}$ of 0 by default for visualization purposes.

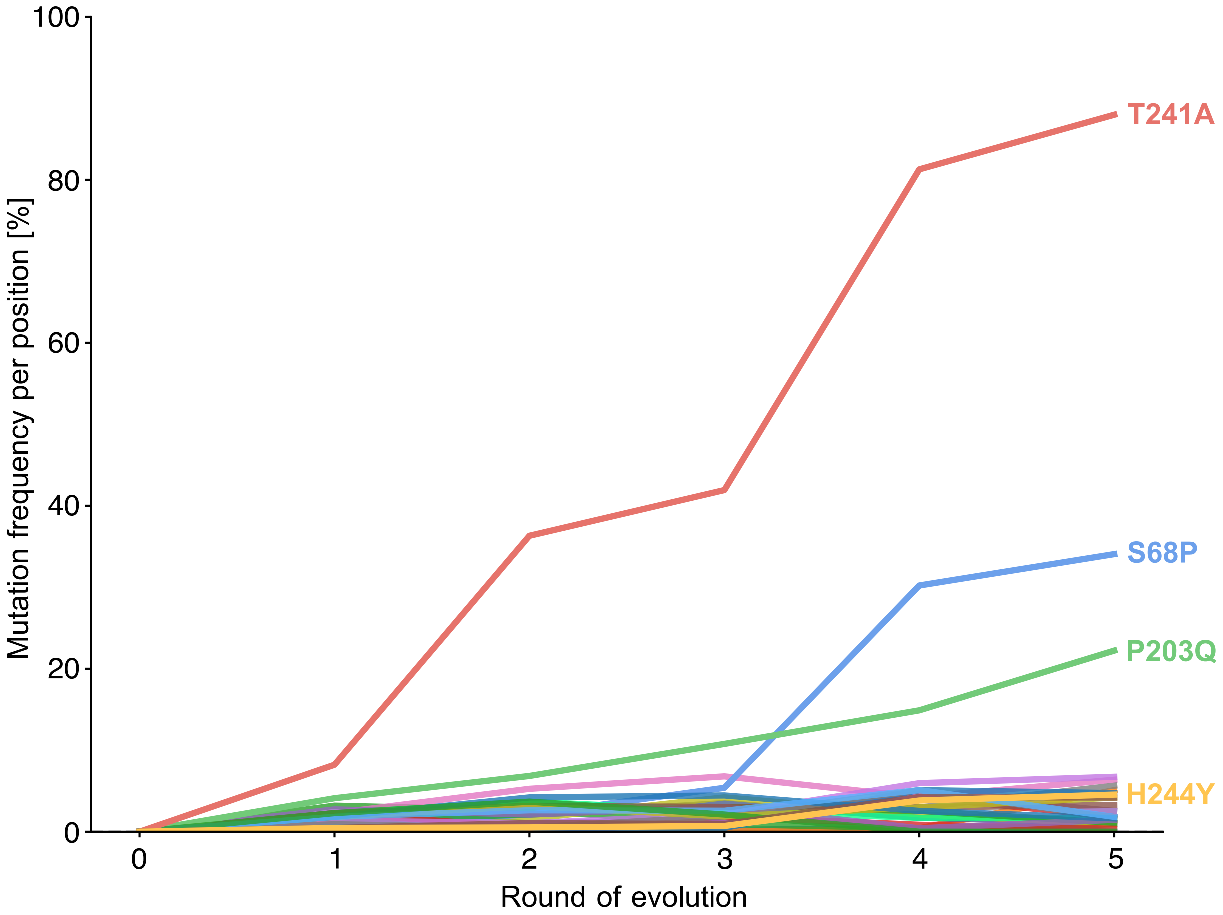

**Figure S12: Mutation frequency of the most mutated positions.** Time-resolved frequency at which the 26 most mutated positions in *Sr*IRED undergo mutation across the entire assayed population. Only the three positions T241A, S68P, and P203Q have significantly spread through the population. The remaining 23 positions (5, 13, 24, 33, 34, 36, 37, 38, 61, 72, 96, 97, 103, 120, 143, 175, 191, 197, 210, 216, 217, 244, 267) are fluctuating between being 0% and 7% mutated.

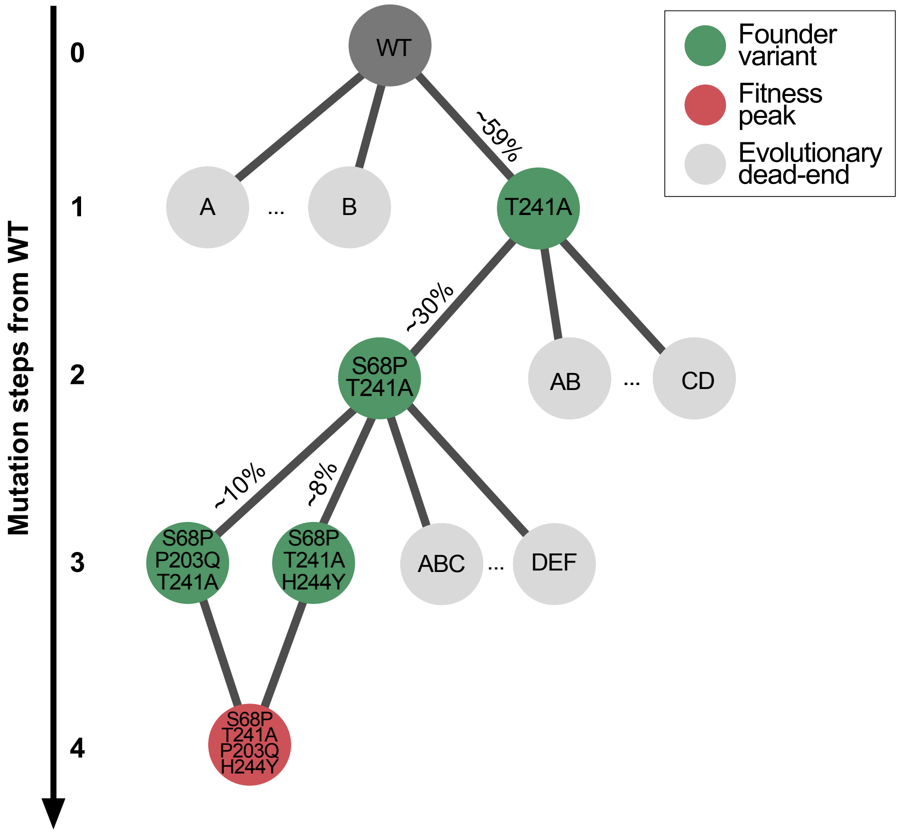

**Figure S13: Phylogeny of founder variants.** Founder variants were identified from the evolution campaign and number of trajectories which pass through each founding node were quantified and displayed in percentage.

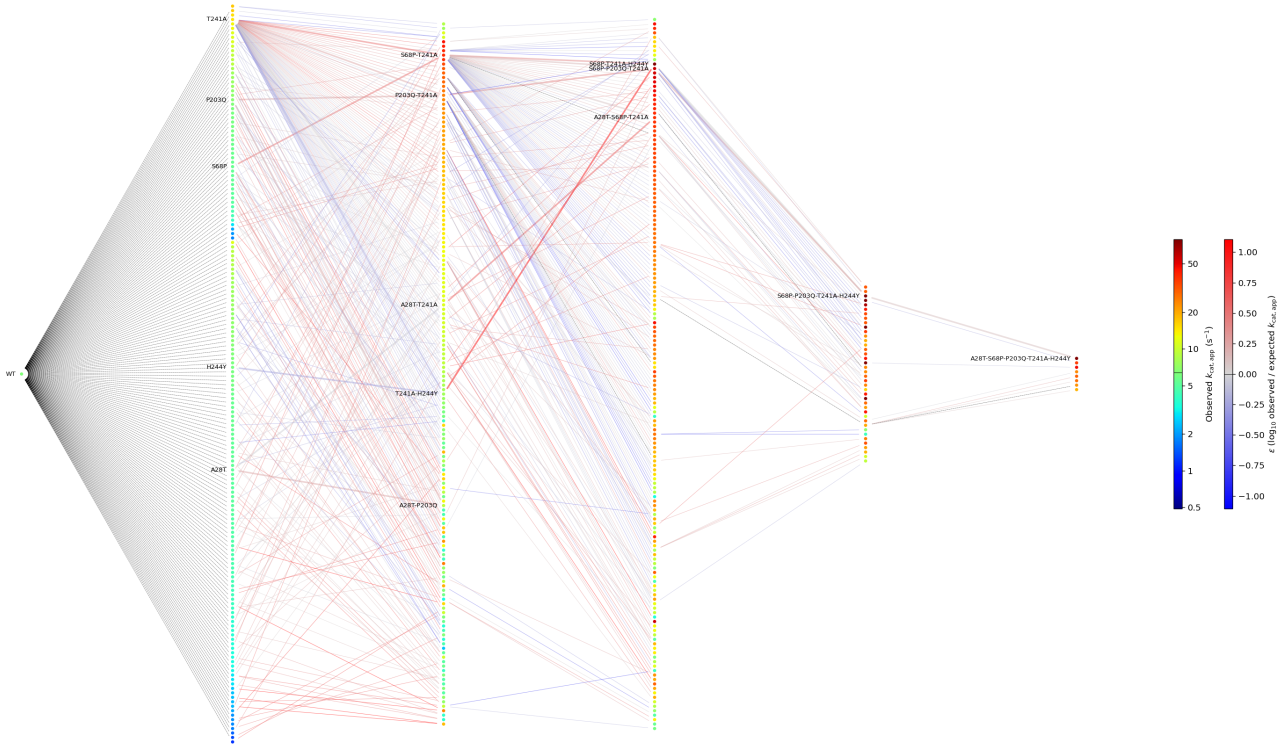

**Figure S14: Connectogram covering the mutants explored in this study with their respective epistatic effects**. Each connected and measured mutational step is coloured by the magnitude of the deviation from an additive expectation in $k_{cat}^{app}$.

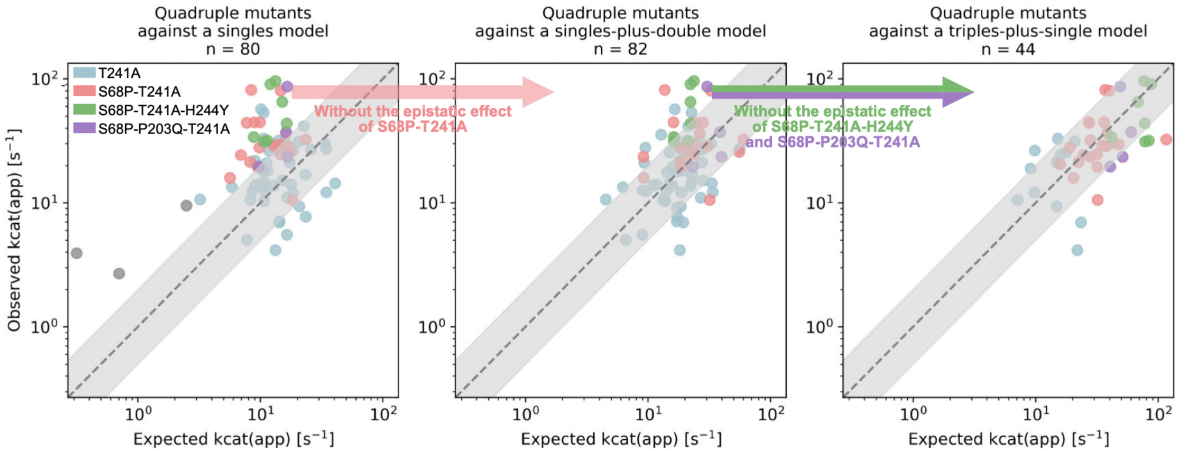

**Figure S15: Analysis of epistasis in quadruple mutants.** A high proportion of highly active quadruple mutants is positively epistatic (left panel) if expected fitness is calculated from single mutation data. Once double mutation data (i.e. the positive epistasis effect of S68P-T241A) or triple mutant data (i.e. the positive epistasis effect of S68P-P203Q-T241A or S68P-T241A-H244Y) is taken into account, most additivity can be restored (middle and right panel).

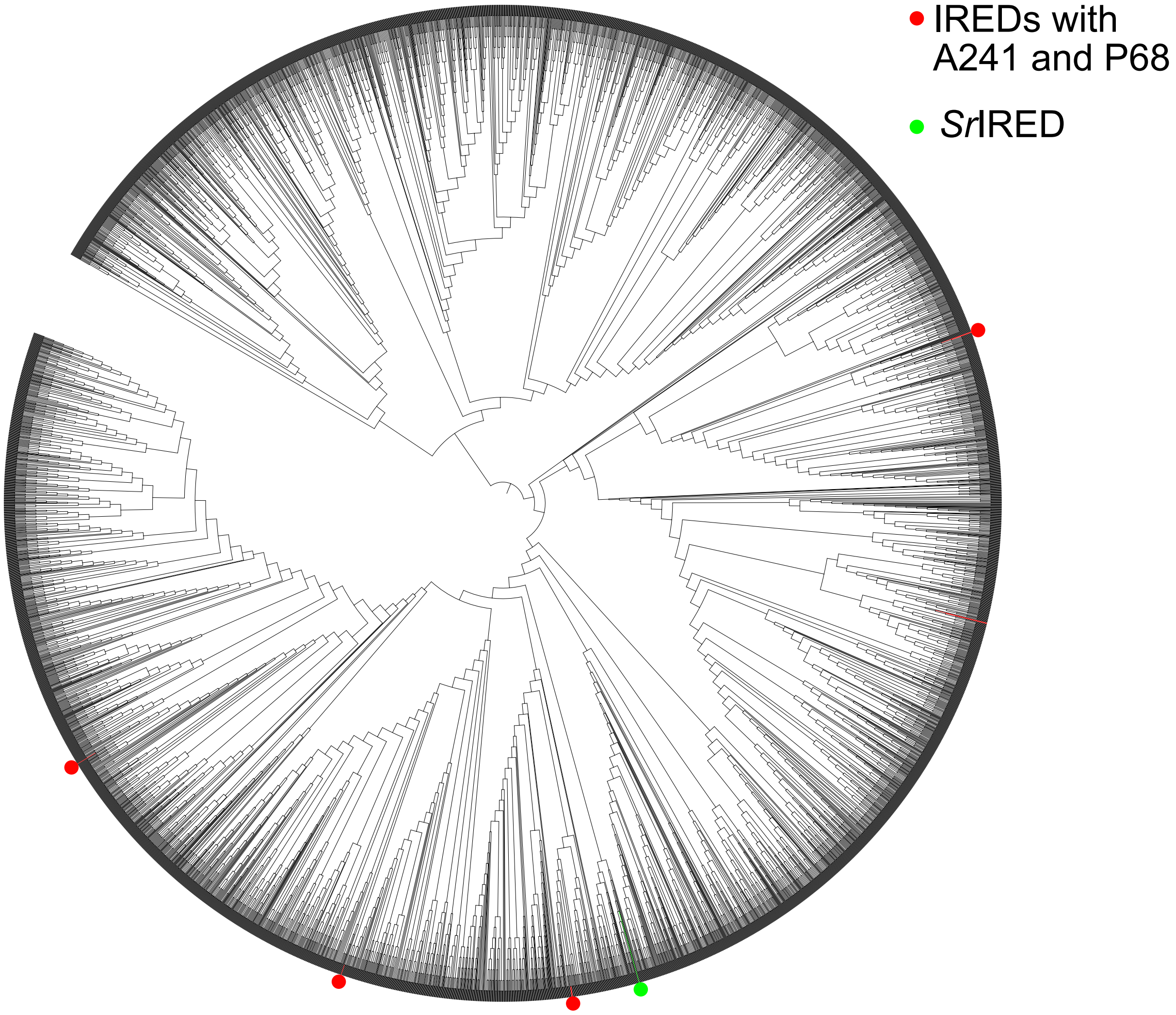

**Figure S16: Phylogenetic tree of 5,000 SrIRED homologues.** The closest 5,000 homologues of *Sr*IRED were retrieved through a BLAST search and a phylogenetic tree was generated using FASTTREE and visualized as a cladogram using iTOL. Indicated are *Sr*IRED (green) and the homologues which contain an alanine at position 241 and a proline at position 68 (red).

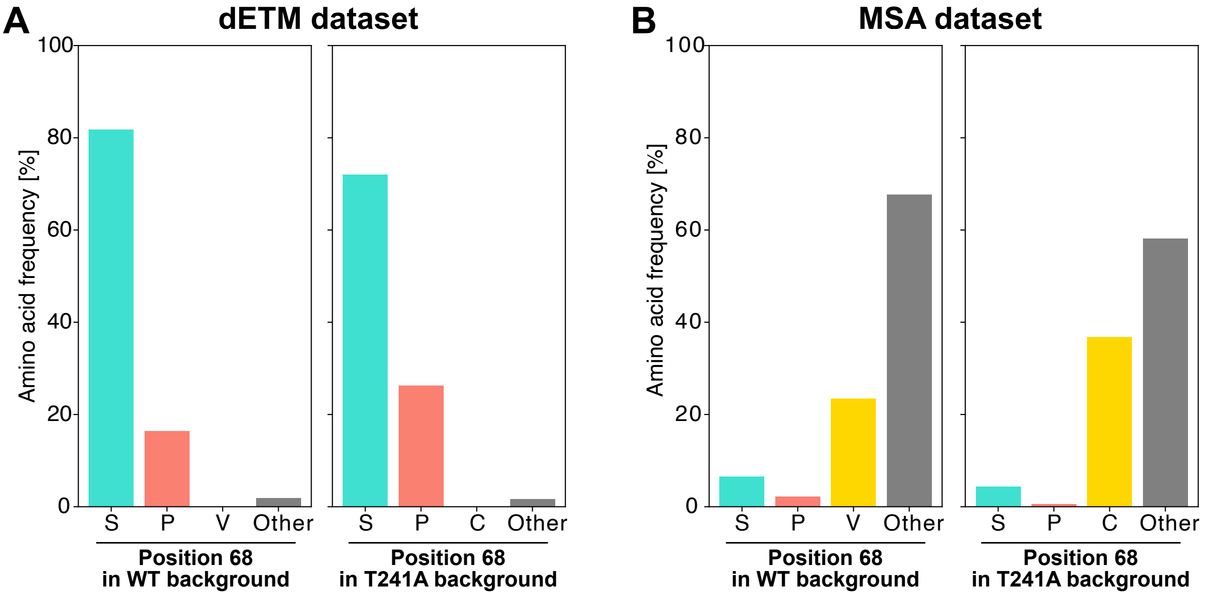

**Figure S17: Proline at position 68 in different mutational backgrounds. (A)** The frequency of a proline at position 68 was assessed in the *Sr*IRED wildtype and *Sr*IRED T241A background. Compared to wildtype, S68P was enriched in the T241A background. **(B)** Frequency of a proline at position 68 in the entire MSA and in the MSA subset which also contained an alanine at position 241. As co-occurrence with A241, proline at position 68 de-enriched in the MSA dataset.

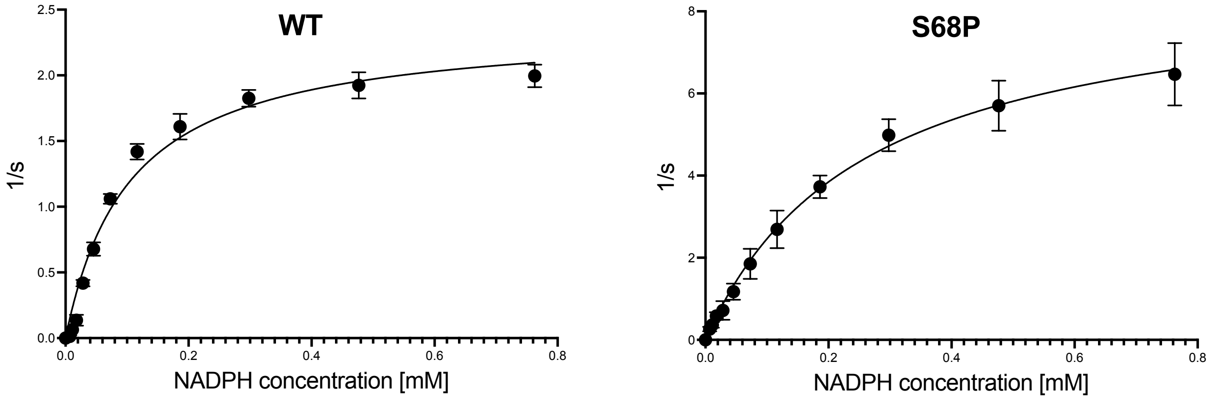

| Mutant | $k_{cat}$ [$s^{-1}$] | $K_M$ [mM] | kcat / $K_M$ [$s^{-1}$ $mM^{-1}$] |
|---|---|---|---|
| Wildtype (WT) | 2.4 ± 0.1 | 0.10 ± 0.01 | 22.8 ± 2.4 |
| S68P | 8.8 ± 0.4 | 0.26 ± 0.03 | 34.1 ± 3.9 |

**Figure S18: NADPH kinetics.** Michaelis-Menten kinetics were conducted with variable concentrations of NADPH (0.007 - 0.763 mM) and a constant concentration of cyclohexanone (30 mM) and cyclopropylamine (30 mM) in Tris pH 8 with variable enzyme concentrations (from 0.004 to 0.01 g/L) depending on the rate at T = 25°C. Initial rates were calculated based on NADPH depletion at 340 nm.

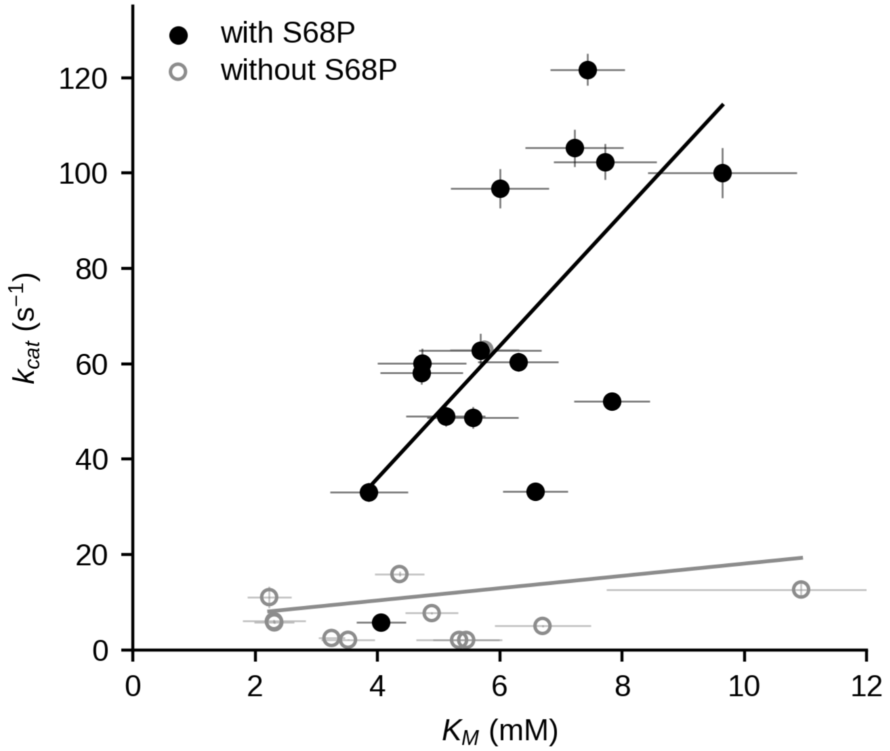

**Figure S19: Trade-off between $k_{cat}$ and $K_M$.** Variants with biophysical measurements from S7 show strong trade-off between the catalytic rate and the substrate (CHX) affinity for variants with S68P (n = 15, r = 0.68, $p$ = 0.005) and without (n = 12, r = 0.19, $p$ = 0.56). This trade-off is driven by the presence of S68P (black) (ANCOVA interaction p = 0.011). Error bars indicate standard deviation (n=3).

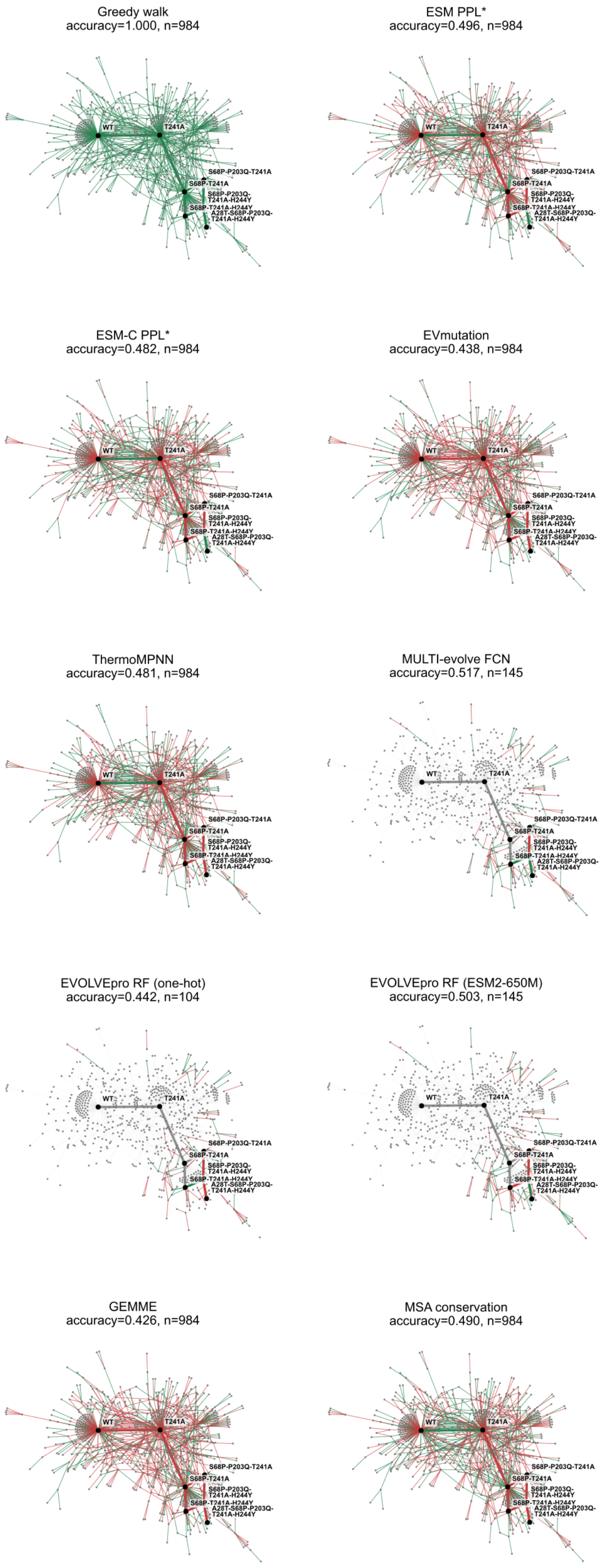

**Figure S20:** Models' ability to predict direction of $k_{cat}^{app}$ change between variants connected in the network graph from the campaign. Steps in which the direction of change was predicted correctly are coloured with green edges, and incorrect predictions are coloured with red edges. The model's accuracy denotes the proportion of correct predictions from the set of predicted edges (n). For supervised models, only the test set of edges are shown.

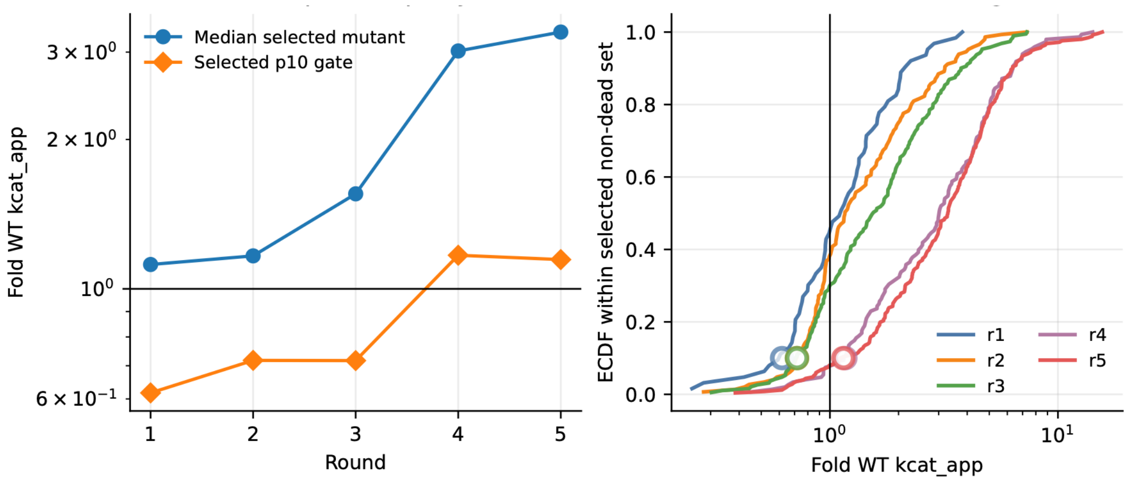

**Figure S21: Quantifying permissive selection over the directed evolution campaign** Selection of variants over the directed evolution campaign analysed (left) by (blue) median $k_{cat}^{app}$ and (orange) the bottom 10th percentile of $k_{cat}^{app}$ in the campaign. Over rounds 1-3, this bottom 10th percentile selected was less fit than the wild-type *Sr*IRED. (right) Empirical Cumulative Distribution Function (ECDF) of the $k_{cat}^{app}$ of variants in each round of the campaign. The bottom 10th percentile in each round is denoted by an open circle, representing a lower threshold for selection in the round.

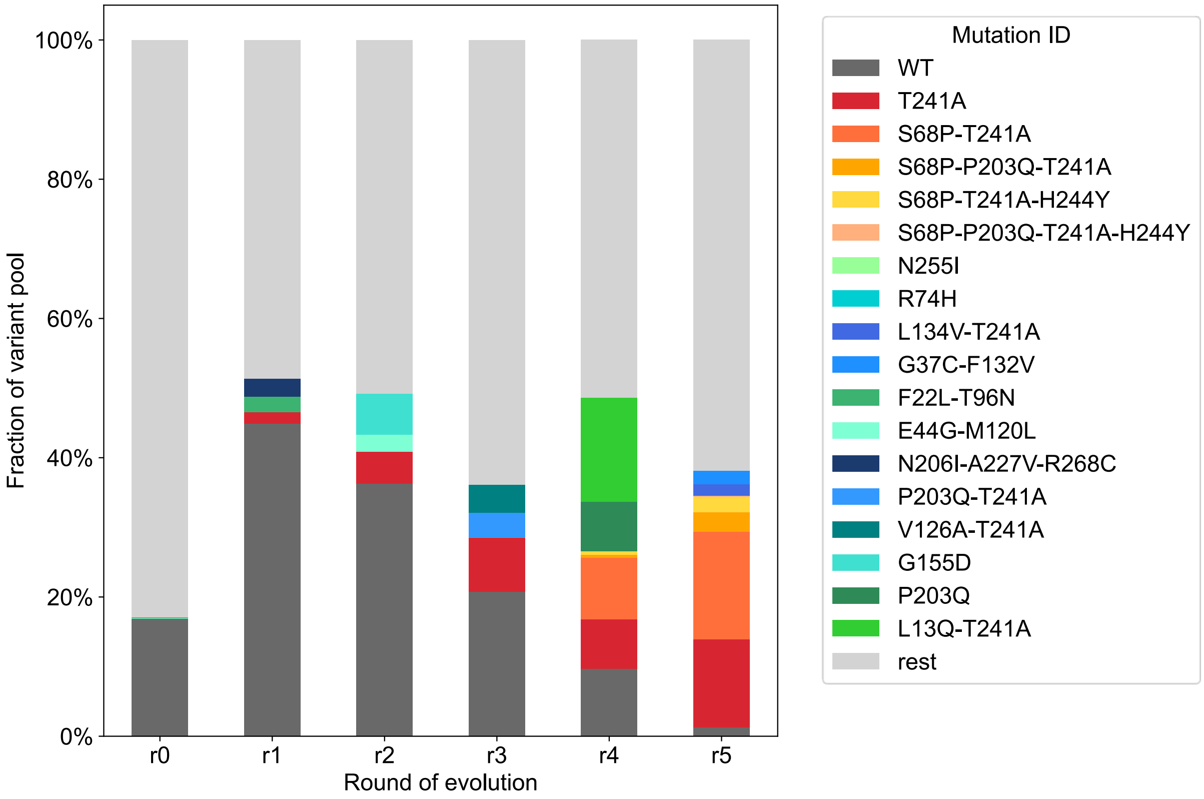

**Figure S22: Fraction of mutants within the variant pool across rounds.** Bar plot with genotype abundance of the founder variants and additional top two most occurring variants in the library in each round. Fraction of variant pool is defined by a mutant's individual number of NGS counts over the total number of counts. Calculated values represent the library proportion each variant takes up after each screening round. Round 0 is the naive 60,000 membered epPCR library before the first round of selection.

**Table S1: 36 hits generated through five rounds of directed evolution.** Hits were defined as variants with more than four-fold improvement in $k_{cat}^{app}$ over wildtype by taking standard deviations into account (hit – std > 4x(wildtype + std)). Trajectories which connected wildtype in single mutation steps to the identified hits were calculated with a total of 144 trajectories being captured.

| Round of evolution | Variant | $k_{cat}^{app}$ | $k_{cat}^{app}$ error | Trajectories to WT |
|---|---|---|---|---|
| r5 | A28T-S68P-P203Q-T241A-H244Y | 100.189 | 29.818 | 9 |
| r5 | S68P-P203Q-T241A-H244Y | 95.954 | 39.588 | 9 |
| r4 | S68P-T241A-H244Y | 90.924 | 13.844 | 3 |
| r5 | S68P-T241A-H244Y-T256M | 90.526 | 35.453 | 3 |
| r5 | S68P-L191V-P203Q-T241A | 86.553 | 22.170 | 10 |
| r4 | S68P-G175D-P203S-T241A | 81.213 | 28.330 | 8 |
| r4 | S68P-E197D-G214R-T241A | 79.872 | 30.475 | 6 |
| r5 | S68P-G175D-P203Q-G221D-T241A | 66.741 | 17.825 | 0 |
| r5 | S68P-L191V-T241A-H244Y | 65.150 | 17.017 | 7 |
| r4 | S68P-P203Q-T241A | 56.885 | 8.441 | 6 |
| r4 | A72T-T241A-H244Y-T256M | 56.697 | 19.616 | 0 |
| r5 | G37C-H238Q-T241A | 56.414 | 11.724 | 3 |
| r5 | S68P-E143D-T241A | 54.497 | 22.196 | 4 |
| r5 | M1V-S68P-T241A | 53.731 | 17.884 | 2 |
| r4 | A72V-P203Q-T241A-G253S | 53.501 | 10.385 | 0 |
| r4 | S68P-E210G-T241A | 50.742 | 12.742 | 6 |
| r5 | S68P-T241A-M282I | 48.624 | 16.444 | 3 |
| r5 | S68P-D69N-G175D-P203S-T241A | 47.341 | 14.934 | 8 |
| r3 | N33S-S51G-T241A | 46.414 | 13.408 | 1 |
| r5 | N33S-T241A | 45.635 | 8.124 | 1 |
| r4 | R54W-S68P-L181M-T241A | 44.743 | 8.908 | 3 |
| r4 | S68P-A194V-T241A-A271E | 44.580 | 8.571 | 4 |
| r4 | S68P-V216I-T241A | 44.496 | 8.164 | 4 |
| r5 | S68P-P203S-T241A-H244Y | 43.727 | 9.397 | 12 |
| r5 | F184L-V216I-T241A | 42.231 | 9.173 | 3 |
| r4 | S68P-L181M-T241A | 41.966 | 9.289 | 3 |
| r5 | S68P-G175D-T241A | 41.912 | 7.721 | 2 |
| r4 | S68P-A205V-T241A | 40.605 | 9.401 | 3 |
| r3 | R34H-T241A | 40.576 | 7.137 | 2 |
| r3 | S68P-T241A | 39.891 | 5.057 | 2 |
| r5 | V76M-T103M-D130E-G226R-T241A | 39.874 | 8.815 | 0 |
| r4 | S68P-G192D-T241A | 39.592 | 6.820 | 4 |
| r3 | V67A-T241A | 39.512 | 8.483 | 1 |
| r5 | S68P-L231Q-T241A | 38.358 | 7.913 | 2 |
| r5 | S68P-D224G-T241A | 37.524 | 7.632 | 2 |
| r5 | S68P-A194V-P203S-T241A | 36.248 | 7.028 | 8 |

**Table S2: Melting temperatures and cyclohexanone kinetics.** Melting temperatures were measured in thermal shift assays in three replicates and expressed as change in melting temperature compared to wildtype ($\Delta T_M$). Errors represent standard deviations. Absolute $T_M$ value of wildtype was 38.5 ± 0.6 °C. Michaelis-Menten kinetics were conducted with variable concentrations of cyclohexanone (0.1 - 52.7 mM) and a constant concentration of cyclopropylamine (75 mM) and NAPDH (1.5 mM) in Tris pH 8 with variable enzyme concentrations (from 0.0015 to 0.05 g/L) depending on the rate at T = 25°C. Initial rates were calculated based on NADPH depletion at 380 nm using a standard curve of NADPH at that wavelength. Fold-change of wildtype is based on $k_{cat}$ measurements. All raw Michaelis Menten plots are provided in **Figure** .

| Mutant | $\Delta T_M$ [°C] | $k_{cat}$ [$s^{-1}$] | $K_M$ [mM] | $k_{cat}$ / $K_M$ [$s^{-1}$ $mM^{-1}$] | Fold-change of wildtype |
|---|---|---|---|---|---|
| Wildtype | - | 4.7 ± 0.1 | 9.1 ± 0.8 | 0.5 ± 0.0 | - |
| T241A | 10.3 ± 0.5 | 8.7 ± 0.2 | 10.6 ± 0.6 | 0.8 ± 0.1 | 2 |
| S68P | -1.4 ± 1.5 | 19.1 ± 0.9 | 19.9 ± 2.1 | 1.0 ± 0.1 | 4 |
| H244Y | -3.4 ± 1.0 | n.d. | n.d. | n.d. | - |
| P203Q | 1.5 ± 0.9 | n.d. | n.d. | n.d. | - |
| G37C | -1.6 ± 1.9 | 7.2 ± 0.2 | 15.7 ± 1.2 | 0.5 ± 0 | 2 |
| A72T | -1.1 ± 0.4 | 8.2 ± 0.5 | 15.1 ± 2.1 | 0.5 ± 0.3 | 2 |
| S68P-T241A | 6.8 ± 1.1 | 106.1 ± 3.2 | 16.7 ± 1.2 | 6.4 ± 0.5 | 23 |
| G37C-T241A | 8.2 ± 1.4 | 17 ± 0.4 | 10.2 ± 0.7 | 1.7 ± 0.1 | 4 |
| A72T-T241A | 9.1 ± 0.4 | 27.9 ± 1.0 | 10.4 ± 1.0 | 2.7 ± 0.3 | 6 |
| S68P-T241A-H244Y | 3.2 ± 2.1 | 188.4 ± 4.5 | 23.1 ± 1.2 | 8.2 ± 0.5 | 40 |
| G37C-S68P-T241A | 4.5 ± 2.7 | 53.7 ± 2.4 | 18.9 ± 2.0 | 2.8 ± 0.3 | 11 |
| S68P-A72T-T241A | 10.6 ± 0.9 | 35.4 ± 1.3 | 15.7 ± 1.5 | 2.3 ± 0.2 | 8 |
| S68P-P203Q-T241A-H244Y | 1.4 ± 1.3 | 205.2 ± 5.5 | 15.4 ± 1.0 | 13.3 ± 0.9 | 44 |

n.d. = not determined

**Table S3: Different bottlenecking strategies for round 2 of the directed evolution campaign**. Using the recorded dETM data, variant T241A was fixed in round 1 and different selection regimes for round 2 were simulated by selecting the rank 1, 10, 12, 13, and 31 mutants of that round. Other mutants below rank 13 disqualified for selection because they contained more than two amino acid changes, their phylogeny died out before round 5, or they encoded for the same phylogenies. For round 3, 4, and 5, the descendant with highest fitness and a maximum of two amino acid changes was chosen.

| **Direct trajectory to the fitness peak of the campaign** | | | | |
|---|---|---|---|---|
| **Round** | **Variant** | **Fitness** | **Sigma** | **Rank** |
| 1 | T241A | 1.3 | 0.2 | 6 |
| 2 | T241A | 1.3 | 0.2 | 31 |
| 3 | S68P-T241A | 2.9 | 0.2 | 5 |
| 4 | S68P-T241A-H244Y | 4.2 | 0.2 | 1 |
| 5 | S68P-P203Q-T241A-H244Y | 4.3 | 0.6 | 2 |
| **Alternative path** | | | | |
| **Round** | **Variant** | **Fitness** | **Sigma** | **Rank** |
| 1 | T241A | 1.3 | 0.2 | 6 |
| 2 | K38N-E210G-T241A | 3.2 | 0.6 | 1 |
| 3 | K38N-E210G-T241A | 3.2 | 0.6 | 1 |
| 4 | K38N-P203Q-T241A | 2.6 | 0.3 | 30 |
| 5 | K38N-T241A-T264M | 2.7 | 0.4 | 44 |

**Table S4: Abundance of winning trajectory variants in the input pool of Round 5**. The number of colonies (*C*) needed to be screened for a given confidence (*P*) of discovering the given variant is determined by its pool abundance (A): $C = \frac{\ln(1-P)}{\ln(1-A)}$ – this determines how many plates need to be screened to discover the hit ($C/96$).

| Variant | Abundance (%) | Clones to screen for 95% confidence of picking | Plates to screen for 95% confidence of picking | Clones to screen for 99% confidence of picking | Plates to screen for 99% confidence of picking |
|---|---|---|---|---|---|
| T241A | 12.5934 | 23 | 1 | 35 | 1 |
| S68P-T241A | 15.4412 | 18 | 1 | 28 | 1 |
| S68P-P203Q-T241A | 2.8567 | 104 | 2 | 159 | 2 |
| S68P-T241A-H244Y | 2.2146 | 134 | 2 | 206 | 3 |
| S68P-P203Q-T241A-H244Y | 0.1168 | 2565 | 27 | 3942 | 42 |
